# Prime-boost vaccination of mice and Rhesus macaques with two novel adenovirus vectored COVID-19 vaccine candidates

**DOI:** 10.1101/2020.09.28.311480

**Authors:** Shengxue Luo, Panli Zhang, Bochao Liu, Chan Yang, Chaolan Liang, Qi Wang, Ling Zhang, Xi Tang, Jinfeng Li, Shuiping Hou, Jinfeng Zeng, Yongshui Fu, Jean-Pierre Allain, Tingting Li, Yuming Zhang, Chengyao Li

## Abstract

COVID-19 vaccines are being developed urgently worldwide, among which single-shot adenovirus vectored vaccines represent a major approach. Here, we constructed two novel adenovirus vectored COVID-19 vaccine candidates on simian adenovirus serotype 23 (Sad23L) and human adenovirus serotype 49 vectors (Ad49L) carrying the full-length gene of SARS-CoV-2 spike protein (S), designated Sad23L-nCoV-S and Ad49L-nCoV-S vaccines, respectively. The immunogenicity elicited by these two vaccine strains was individually evaluated in mice. Specific humoral and cellular immune responses were proportionally observed in a dose-dependent manner, and stronger response was obtained by boosting. Furthermore, five rhesus macaques were intramuscularly injected with a dose of 5×10^9^ PFU Sad23L-nCoV-S vaccine for prime vaccination, followed by boosting with 5×10^9^ PFU of Ad49L-nCoV-S vaccine at 4-week interval. Three macaques were injected with Sad23L-GFP and Ad49L-GFP vectorial viruses as negative controls. Both mice and macaques tolerated well the vaccine inoculations without detectable clinical or pathologic changes. In macaques, prime-boost vaccination regimen induced high titers of 10^3.16^ S-binding antibody (S-BAb), 10^2.75^ cell receptor binding domain (RBD)-BAb and 10^2.38^ neutralizing antibody (NAb) to pseudovirus a week after boosting injection, followed by sustained high levels over 10 weeks of observation. Robust IFN-γ secreting T-cell response (712.6 SFCs/10^6^ cells), IL-2 secreting T-cell response (334 SFCs/10^6^ cells) and intracellular IFN-γ expressing CD4^+^/CD8^+^ T cell response (0.39%/0.55%) to S peptides were detected in the vaccinated macaques. It was concluded that prime-boost immunization with Sad23L-nCoV-S and Ad49L-nCoV-S vaccines can safely elicit strong immunity in animals in preparation of clinical phase 1/2 trials.

## INTRODUCTION

Novel coronavirus disease 2019 (COVID-19) usually presents as severe acute respiratory syndrome triggered by SARS-CoV-2 infection (*1, 2*), which has become globally pandemic and killed nearly one million people worldwide (WHO Coronavirus Disease [COVID-19] Dashboard) (*3*). Currently the most urgent need is to develop safe and effective vaccines that prevent SARS-CoV-2 infection.

SARS-CoV-2 is a positive-sense single-stranded RNA virus, encoding four structural proteins, including spike (S), envelope (E), membrane (M) and nucleocapsid (N) (*4, 5*). The S protein is a glycoprotein carrying the cell receptor binding domain (RBD), and is a major protective antigen that may elicit potent neutralizing antibody (NAb) and cellular immunity (*4, 5*). Therefore, S protein has been the primary target antigen for developing recombinant vaccines.

According to the World Health Organization (WHO) report for a draft landscape of COVID-19 candidate vaccines on August 28, 2020, there are 33 vaccine candidates in clinical evaluation, which are mainly distributed across five types of biotechnological platforms, *i.e.*, inactivated virus, DNA, RNA, protein subunit and non-replicating viral vector vaccines (*6*). Among COVID-19 candidate vaccines in clinical trials, five non-replicating adenovirus vectored vaccines progressed to the frontline, of which one (CanSino Biological Inc./Beijing Institute of Biotechnology) has been approved for emerging use in Chinese military personal, one (Gamaleya Research Institute) has been registered in Russia, and two have been in initial phase III clinical trial (University of Oxford/AstraZeneca; Janssen Pharmaceutical Companies), and one in phase I clinical trial (ReiThera/LEUKOCARE/Univercells), respectively. Recombinant adenovirus vectors originated from various serotype of strains have displayed good safety profiles and induced broad and strong humoral and cellular immune responses, which have been widely used for research and development of vaccines (*7*). Based on the published data regarding human adenovirus type 5 (Ad5), chimpanzee adenovirus type 25Y (ChAdOx1) and human adenovirus type 26 (Ad26) vectorial COVID-19 vaccines (*8–14*), results showed that a single-shot vaccine prevented SARS-CoV-2 pneumonia in rhesus macaques and hamsters, and elicited significant immune response in the majority of recipients in phase I/II clinical trials. However, relatively weaker protection in animals and lower immune response in humans were observed with a single dose of vaccine compared with prime-boost immunizations by two or three doses of inactivated virus or mRNA vaccines (*15–19*). Enhancement of immune response was evidenced by prime and boost vaccination regimens with homologous boosting of ChAdOx1 nCoV-19 in animals and humans (*11, 20*), or with heterologous Ad26 and Ad5 vectored COVID-19 vaccines in humans (*21*).

In this study, two novel simian adenovirus type 23 (SAdV23) and human adenovirus type 49 (HAdV49) derived vectors (Sad23L and Ad49L) were used for developing the COVID-19 vaccines carrying the full-length S gene of SARS-CoV-2 (*22, 23*) designated Sad23L-nCoV-S and Ad49L-nCoV-S vaccines, respectively. These two adenovirus vectored COVID-19 vaccines presented high infectious titers and low frequencies of pre-existing immunity in humans. Immunogenicity was extensively evaluated in mice and rhesus macaques by prime-boost vaccinations with these two novel heterologous adenovirus vectored vaccines.

## RESULTS

### Production and characterization of Sad23L-nCoV-S and Ad49L-nCoV-S vaccines

An optimized and synthesized full-length S gene of SARS-CoV-2 was cloned into the deleted E1 region under cytomegalovirus (CMV) promotor regulation within two novel adenovirus vectorial Sad23L and Ad49L plasmids, designated as Sad23L-nCoV-S or Ad49L-nCoV-S, respectively (Fig. 1A). The recombinant adenoviruses were rescued from HEK-293A. A large amount of Sad23L-nCoV-S and Ad49L-nCoV-S candidate vaccines were produced from HEK-293A cell cultures, and further purified and titrated over 10^11^ PFU/ml.

**Fig 1.**
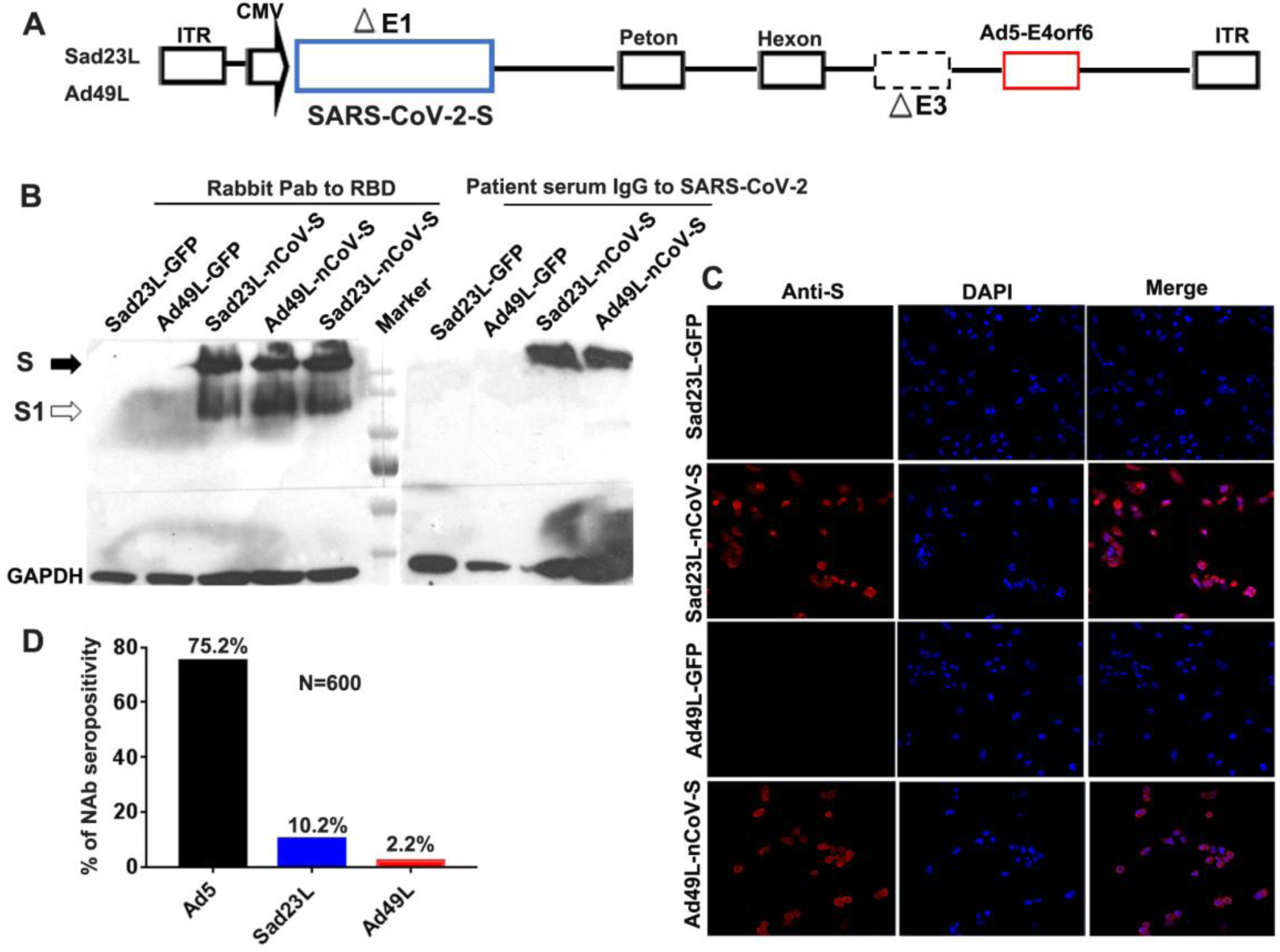
Characterization of Sad23L-nCoV-S and Ad49L-nCoV-S vaccines. (**A**) Recombinant adenovirus constructs Sad23L-nCoV-S and Ad49L-nCoV-S carrying the full-length S gene of SARS-CoV-2 under CMV promotor regulation within the deleted E1 region of Sad23L or Ad49L vector. (**B**) Western blot analysis for the expression of S protein from Sad23L-nCoV-S or Ad49L-nCoV-S infected HEK-293A cell lysates by rabbit polyclonal antibody to RBD and heat-inactivated COVID-19 patent’s serum IgG. Sad23L-GFP or Ad49L-GFP virus infected cells were used as mock controls. (**C**) Expression of S protein in HEK-293A cells detected by immunofluorescence staining. (**D**) Seroprevalence of neutralizing antibody to Ad5, Ad49L or Sad23L vector in 600 healthy blood donors.

Expression of S protein was verified in Sad23L-nCoV-S and Ad49L-nCoV-S vaccine strains infected HEK-293A cells by Western blot with rabbit polyclonal anti-RBD antibodies and COVID-19 patient’s serum IgG, respectively, but not in the Sad23L-GFP and Ad49L-GFP vectorial viruses infected cells (Fig. 1B). The expression of S protein in the vaccine strains-infected HEK-293A cells was also observed in red color by an immunofluorescence staining, but not in the adenovirus-GFP infected control cells (Fig. 1C). These results indicated that Sad23L-nCoV-S and Ad49L-nCoV-S vaccines could effectively produce SARS-CoV-2 S protein in the infected cells.

To measure pre-existing immunity to these two vectors, 600 healthy blood donor samples were collected from six cities crossing the south, north, east, west and central regions of China and were tested for neutralizing antibodies (NAb) reacting with Sad23L-GFP, Ad49L-GFP and Ad5-GFP viruses, respectively. The seroprevalence of Sad23L, Ad49L and Ad5 was 10.2%, 2.2% or 75.2% respectively (Fig. 1D), indicating that Sad23L and Ad49L vectors had low pre-exposure rate in the Chinese population.

### Animal tolerance of Sad23L-nCoV-S and Ad49L-nCoV-S vaccine inoculation

C57BL/6 mice (n=5 per group) were intramuscularly injected with individual Sad23L-nCoV-S and Ad49L-nCoV-S vaccines at doses of 10^7^, 10^8^ and 10^9^ PFU or by prime-boost immunization with these two adenovirus vectored vaccines (10^9^ PFU/each) at 4 week interval and compared with vectors and PBS controls, respectively. Mice tolerated well the various doses of vaccines or vector controls at 28 or 56 days, presenting no obvious change of weight and body temperature when compared with injected PBS (Fig. 2A). In addition, histopathological examination of brain, lung, heart, liver, kidney and muscle tissues (at intramuscular injection site and para-tissues) did not present significant pathological lesions over 4 weeks following the prime only or prime-boost injections (Fig. S1).

**Fig. 2.**
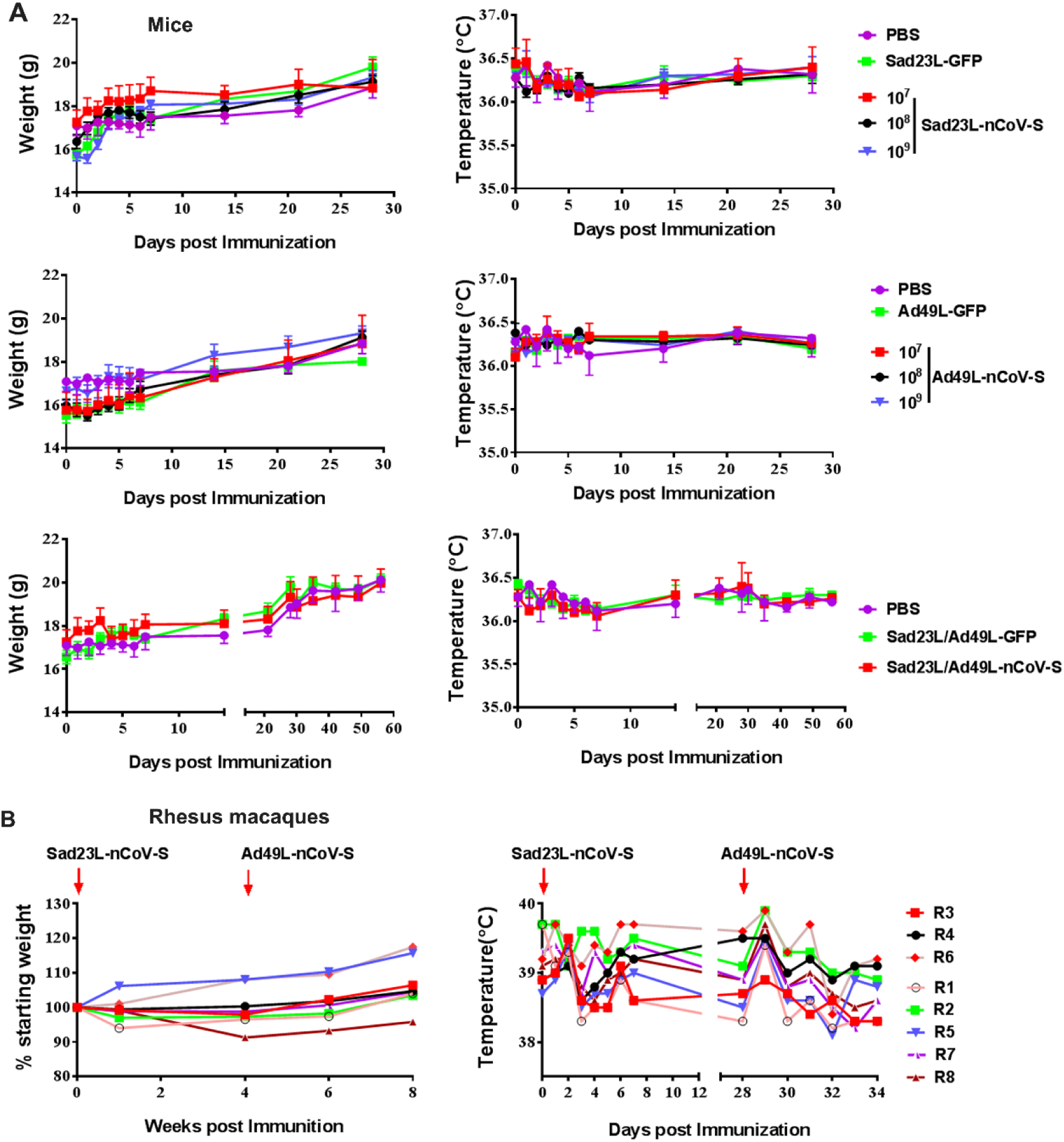
Examination of weight and body temperature of mice and rhesus macaques inoculated with Sad23L-nCoV-S and Ad49L-nCoV-S vaccines. Animals were intramuscularly immunized by prime only with either Sad23L-nCoV-S or Ad49L-nCoV-S vaccine at three different doses, or by prime-boost with first Sad23L-nCoV-S and boost with Ad49L-nCoV-S vaccines at 4 week interval. Body weight and temperature were monitored during the course up to 60 days for mice or 8 weeks for monkeys. (**A**) C57BL/6 mice (n=5/group) immunized with a dose of 10^7^, 10^8^ or 10^9^ PFU Sad23L-nCoV-S vaccine, 10^9^ PFU Sad23L-GFP and an equal volume of PBS controls (top panel); a dose of 10^7^, 10^8^ or 10^9^ PFU Ad49L-nCoV-S vaccine, 10^9^ PFU Ad49L-GFP and an equal volume of PBS controls (middle panel); a dose of 10^9^ PFU Sad23L-nCoV-S followed by a dose of 10^9^ PFU Ad49L-nCoV-S, or 10^9^ PFU Sad23L-GFP and 10^9^ PFU Ad49L-GFP vectorial viruses and an equal volume of PBS controls (low panel). (**B**) Rhesus monkeys (11-14y) inoculated with a dose of 5×10^9^ PFU Sad23L-nCoV-S and a dose of 5×10^9^ PFU Ad49L-nCoV-S vaccines (R1, R2, R5, R7 and R8), or 5×10^9^ PFU Sad23L-GFP and a dose of 5×10^9^ PFU Ad49L-GFP controls (R3, R4 and R6).

Vaccination (n=5) and sham control (n=3) groups of rhesus macaques (Table S1) were first injected with 5×10^9^ PFU Sad23L-nCoV-S vaccine or Sad23L-GFP control, and then at 4 week interval, animals were injected with a second dose of 5×10^9^ PFU Ad49L-nCoV-S vaccine or Ad49L-GFP control, respectively. During the course of immunization, clinical parameters were monitored for eight weeks. All monkeys displayed normal appetite and mental state, and no obvious change of weight and body temperature was observed with these eight animals (Fig. 2B). Hematological and biochemical examination of blood samples showed no notable variation when comparing the vaccinated or vector control animals during 8 weeks pre-vaccination and post-vaccination (Fig. S2; Table S2).

### A single-shot immunization of Sad23L-nCoV-S or Ad49L-nCoV-S vaccine induced specific immune response in mice

To evaluate the immunogenicity of individual Sad23L-nCoV-S and Ad49L-nCoV-S vaccines, a single dose of 10^7^, 10^8^ or 10^9^ PFU Sad23L-nCoV-S or Ad49L-nCoV-S vaccine was intramuscularly injected to C57BL/6 mice (n=5/group). Vector control groups (n=5/group) received 10^9^ PFU Sad23L-GFP or Ad49L-GFP viruses, and naïve control group (n=5) received an equal volume of PBS, respectively. Four weeks post-immunization, titers of S or RBD binding antibody (S-BAb or RBD-BAb) were quantified by ELISA in a dose-dependent manner with both Sad23L-nCoV-S and Ad49L-nCoV-S immunized mice, but not in the control groups (*P*<0.001, Fig. 3, A-D). A dose of 10^9^ PFU Sad23L-nCoV-S vaccine induced antibody titers of 10^4.27^ S-BAb and 10^3.98^ RBD-BAb (Fig. 3, A and B), while the same dose of Ad49L-nCoV-S vaccine induced titers of 10^3.02^ S-BAb and 10^2.42^ RBD-BAb, respectively (Fig. 3, C and D).

**Fig 3.**
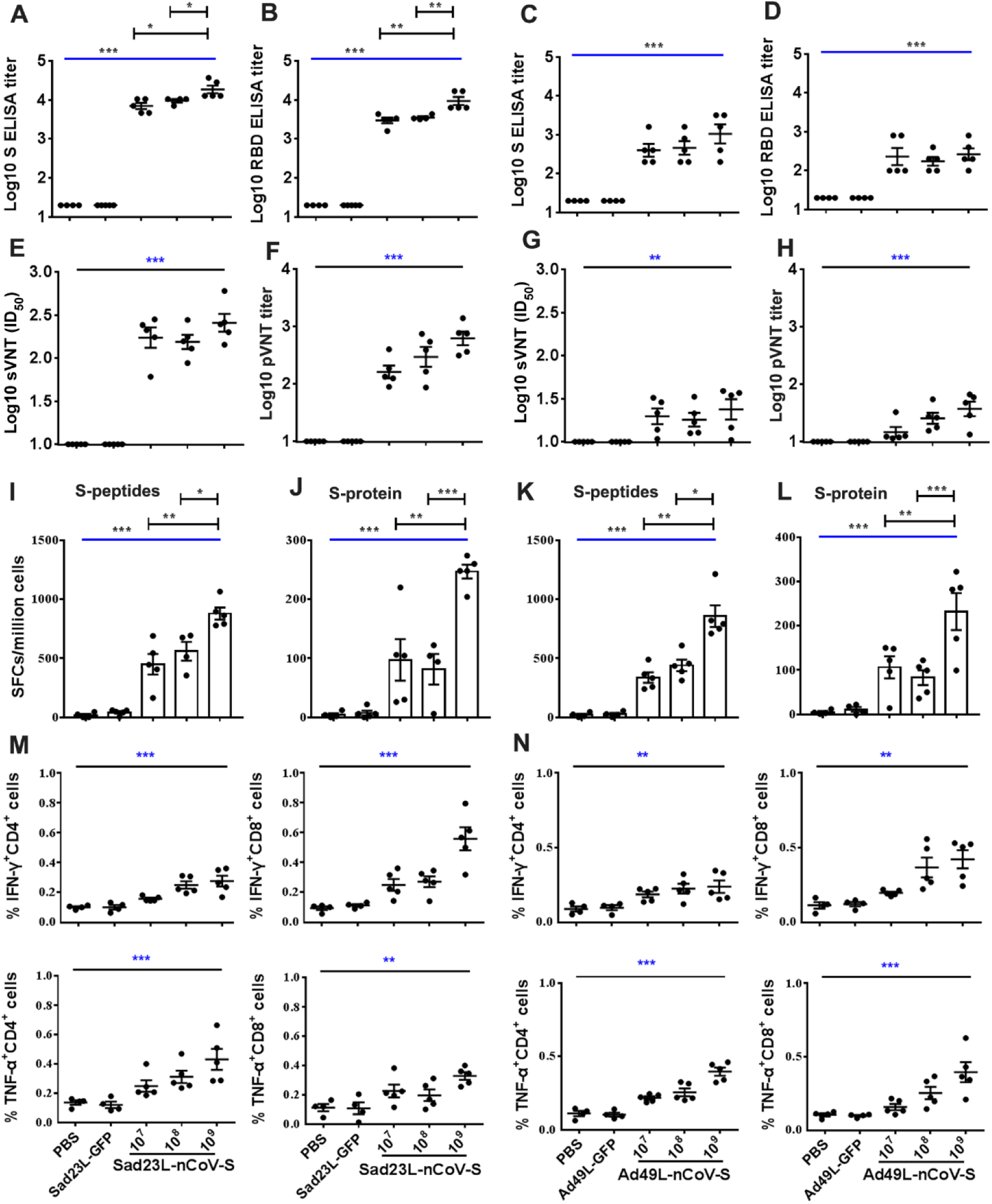
Specific antibody and T cell response of C57BL/6 mice inoculated with a single shot of Sad23L-nCoV-S or Ad49L-nCoV-S vaccine at three different doses. C57BL/6 mice (n=5/group) were immunized by a single dose of 10^7^, 10^8^ or 10^9^ PFU Sad23L-nCoV-S or Ad49L-nCoV-S vaccine. Mice sera and splenocytes were collected for measurement of antibody level and T cell response 4 weeks post-immunization. (**A-D**) S or RBD binding antibody (S-BAb or RBD-BAb) titers were obtained by ELISA. (**E-H**) Neutralizing antibody (NAb) titers were obtained by surrogate virus-based neutralization test (sVNT) or pseudovirus-based neutralization test (pVNT). (**I-L**) IFN-γ secreting T cell response (spot forming cells [SFCs]/million cells) of splenocytes to S peptides and S protein from Sad23L-nCoV-S or Ad49L-nCoV-S immunized mice was measured by ELISpot, respectively. (**M** and **N**) Frequency of IFN-γ or TNF-α expressing CD4^+^ and CD8^+^ T cell response of splenocytes to S peptides from Sad23L-nCoV-S or Ad49L-nCoV-S immunized mice was obtained by ICS. Data is shown as mean ± SEM (standard errors of means). *P* values are analyzed by one-way ANOVA with 2-fold Bonferroni adjustment. Statistically significant differences are shown with asterisks (*, *P*<0.05; **, *P*< 0.01 and ***, *P*< 0.001).

The neutralizing antibody (NAb) titers to SARS-CoV-2 were titrated in two individual vaccine immunized mice by surrogate virus based NAb test (sVNT) and pseudovirus-based NAb test (pVNT) at 50% inhibitory dilution (ID_50_), respectively (Fig. 3, E-H). A dose of 10^9^ PFU Sad23L-nCoV-S vaccine induced NAb titers of 10^2.41^ sVNT(ID_50_) and 10^2.79^ pVNT (ID_50_) (Fig. 3, E and F), and Ad49L-nCoV-S vaccine induced titers of 10^1.38^ sVNT (ID_50_) and 10^1.57^ pVNT(ID_50_), respectively (Fig. 3, G and H).

Specific T-cell response of isolated splenocytes was examined in vaccinated and sham mice after stimulation with S peptides, S protein and RBD protein, respectively (Fig. 3, I-N; Fig. S3). A single dose of Sad23L-nCoV-S vaccine elicited strong specific IFN-γ secreting T cell response to S peptides (450.2-898.4 SFCs/million cells, Fig. 3I) and S protein (81.5-246.8 SFCs/million cells, Fig. 3J), while a dose of Ad49L-nCoV-S vaccine induced similar pattern of IFN-γ secreting T cell response (336.8-857.1 or 82.8-232 SFCs/million cells, Fig. 3, K and L). Both vaccines elicited T cell response significantly higher than sham controls (*P*<0.001), but response to RBD protein was not significantly different (*P*>0.05, Fig. S3, A and B). Specific intracellular levels of IFN-γ, TNF-α and IL-2 expressing CD4^+^ or CD8^+^ T cell response to S peptides were measured by ICS, in which significantly higher frequency of IFN-γ and TNF-α but not IL-2 expressing CD4^+^/CD8^+^ T-cells was found in both Sad23L-nCoV-S and Ad49L-nCoV-S vaccinated mice compared to sham mice (*P*<0.001, Fig. 3, M and N; Fig. S3, C-F).

Overall, individual Sad23L-nCoV-S or Ad49L-nCoV-S vaccinated mice developed specific BAb and NAb antibodies and T cell responses to S protein, RBD protein or S peptides of SARS-CoV-2 in a dose-dependent fashion (Fig. 3), suggesting strong immunogenicity of the two novel adenovirus vectored COVID-19 vaccines.

### Prime-boost immunization of mice with Sad23L-nCoV-S and Ad49L-nCoV-S vaccines

To further improve reactivity and longevity of immune response to vaccines, the prime-boost vaccination regimen was utilized to immunize C57BL/6 and BALB/c mice (n=5/group) with a prime dose of 10^9^ PFU Sad23L-nCoV-S on day 0 and a boost dose of 10^9^ PFU Ad49L-nCoV-S on day 28 (Fig. 4A). In comparison with prime immunization with Sad23L-nCoV-S, a booster with Ad49L-nCoV-S significantly increased BAb and NAb titers in C57BL/6 and BALB/c mice (*P*<0.001; Fig. 4, B-G), and titers were maintained at high level for 10 weeks of monitoring (Fig. 4, D and G). Profiling of IgG subclasses showed a predominant serum IgG2a to RBD protein associating with Th1 response in prime-boost vaccinated mice (Fig. S4). Regarding specific T cell response, the boosting with Ad49L-sCoV-S on Sad23L-nCoV-S prime immunization enhanced or stabilized specific IFN-γ-secretion T cell responses to S peptides, S protein or RBD protein (Fig. 4H), as well as levels of intracellular cytokines IFN-γ and TNF-α but not of IL-2 response to S peptides compared with single dose of vaccine vaccinated or sham control C57BL/6 and BALB/c mice (Fig. 4I; Fig. S5).

**Fig. 4.**
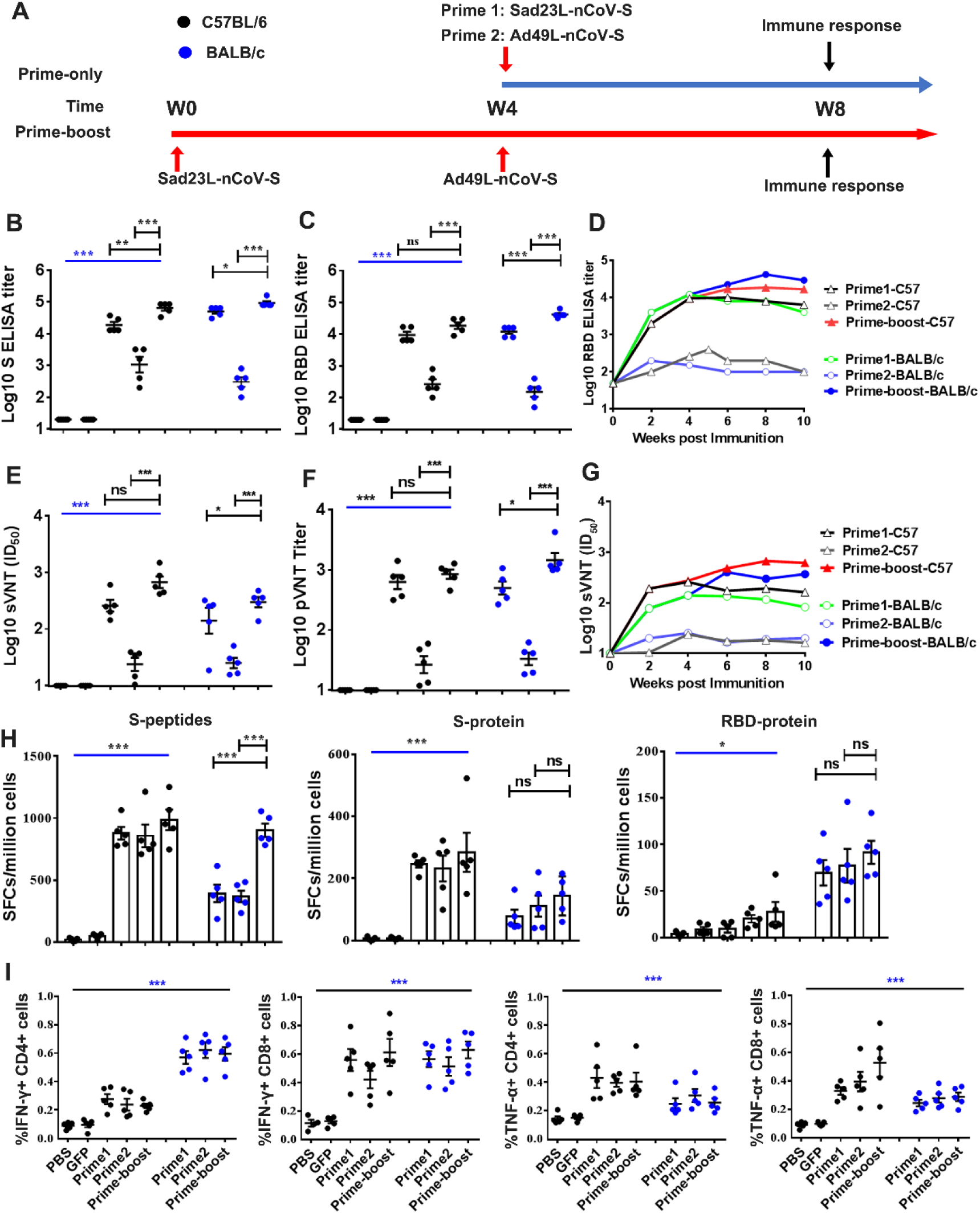
Specific humoral and cellular immune response of C57BL/6 and BALB/c mice to prime-boost immunization with Sad23L-nCoV-S and Ad49L-nCoV-S vaccines. (**A**) C57BL/6 and BALB/c mice (n=5/group) were prime immunized with a dose of 10^9^ PFU Sad23L-nCoV-S vaccine and boosted with a dose of 10^9^ PFU Ad49L-nCoV-S vaccine at 4 week interval. Sera and splenocytes were collected from vaccinated or control mice for measurement of antibody and T cell responses 4 weeks after both prime only and boosting immunizations. (**B-D**) Anti-S-BAb and RBD-Bab titers determined by ELISA. (**E-G**) NAb titers measured by sVNT and pVNT. (**H**) IFN-γ secreting T cell response (SFCs/million cells) to S peptides, S or RBD protein measured by ELISpot. (**I**) Frequency of IFN-γ or TNF-α expressing CD4^+^ and CD8^+^ T cell response to S peptides determined by ICS. Data are shown as a mean ± SEM. *P* values are analyzed with two-tailed t test. Statistically significant differences are shown with asterisks (*, *P*<0.05; **, *P*< 0.01 and ***, *P*< 0.001).

Taken together, the results suggest that prime-boost vaccination of two species of mice with Sad23L-nCoV-S followed by Ad49L-nCoV-S enhanced specific immune response to SARS-CoV-2 when compared with prime vaccination only with a single-shot of Sad23L-nCoV-S or Ad49L-nCoV-S vaccine.

### Rhesus macaques’ specific immune response to prime-boost vaccination with combined Sad23L-nCoV-S and Ad49L-nCoV-S vaccines

Eight rhesus macaques aged 11-14 years were selected and tested for the baseline values of antibody and T-cell response to SARS-CoV-2 S from blood samples in pre-vaccination at week -1 or week 0 (Fig. 5A; Table S1). The pre-existing NAb titer to Sad23L, Ad49L or Ad5 was detected < 1:10 in serum samples from all eight animals (Table. S1).

**Fig. 5.**
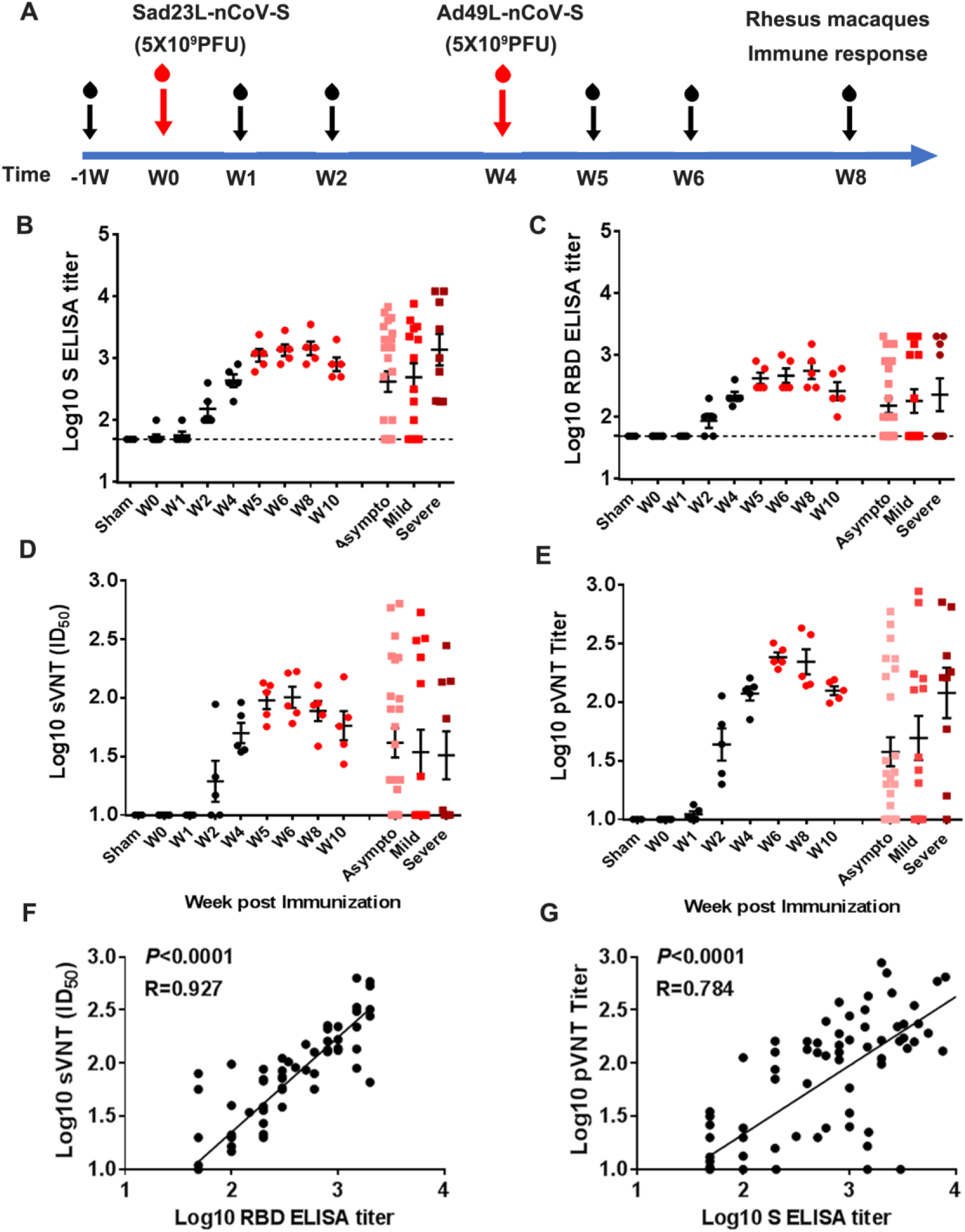
Antibody reactivity of Rhesus macaques to prime-boost vaccination with Sad23L-nCoV-S and Ad49L-nCoV-S vaccines. (**A**) Five rhesus macaques were prime immunized with 5×10^9^ PFU of Sad23L-nCoV-S vaccine and boosted with 5×10^9^ PFU of Ad49L-nCoV-S vaccine at 4 week interval. Blood samples were collected weekly from immunized or sham control macaques. Three macaques first immunized with 5×10^9^ PFU of Sad23L-GFP viruses and boosted with 5×10^9^ PFU of Ad49L-GFP viruses were used as sham controls. Convalescent serum samples from 25 asymptomatic, 14 mild and 9 severe COVID-19 infected patients were taken as positive controls. (**B** and **C**) S-BAb and RBD-BAb titers were tested by ELISA. (**D** and **E**) NAb titers were measured by sVNT and pVNT. (**F** and **G**) Correlation between RBD-BAb and sVNT or S-BAb and pVNT assay were compared using Spearman nonparametric correlation, respectively.

Five rhesus macaques were intramuscularly injected first with a dose of 5×10^9^ PFU Sad23L-nCoV-S vaccine, then with a boost injection of 5×10^9^ PFU Ad49L-nCoV-S vaccine 4 weeks later, while three sham rhesus macaques controls were injected with equal doses of Sad23L-GFP and Ad49L-GFP viruses, respectively (Fig. 5A). Blood samples were collected weekly from these two groups of animals. Both BAb (S-BAb and RBD-BAb) and NAb (sVNT and pVNT) levels were titrated in the vaccinated group but not in sham group. BAb and NAb reactivity increased at week 2 post prime-immunization, at week 4 (W0 for boost-immunization), at weeks 5 and 6 and then stayed at high levels up to week 10 (Fig. 5, B-E). Titers of 10^3.16^ S-BAb (Fig. 5B), 10^2.75^ RBD-BAb (Fig. 5C), 10 ^2.01^ sVNT(ID_50_) (Fig. 5D) and 10^2.38^ pVNT(ID_50_) (Fig. 5E) were quantified and found higher than the corresponding antibody titers found in sera from 25 asymptomatic, 14 mild and 9 severe COVID-19 convalescent patients (Fig. 5, B-E). Strong correlation between BAb and NAb titers was observed (*P*<0.0001, R=0.784-0.927; Fig. 5, F and G), suggesting the reliability of these antibody quantitative assays.

PBMCs were isolated from whole blood of pre- and post-immunized monkeys for evaluation of T cell responses to S peptides, S and RBD protein by ELISpot and ICS (Fig. 6). Prime vaccination with a dose of Sad23L-nCoV-S vaccine induced an increase of IFN-γ secreting T cell response to S peptides (406.6-526.3 SFCs/million cells) at weeks 2 and 4 post prime-vaccination, and then boosting with a dose of Ad49L-nCoV-S vaccines enhanced the IFN-γ secreting T cell response (583.9-712.6 SFCs/million cells) at weeks 5 to 8 (Fig. 6A). IFN-γ secreting T cell reaction to S and RBD proteins stayed at high levels after prime-boost immunizations (Fig. 6, B and C). Relatively high IL-2 secreting T cell response to S peptides, S and RBD proteins was observed (Fig. 6, D-F), but weak IL-4 secreting T cell response by ELISpot (Fig. S6). Frequency of intracellular IFN-γ expressing CD4^+^/CD8^+^ T-cell responses to S peptides was observed in vaccinated macaques at weeks 2 to 8 (Fig. 6, G-J), significantly higher than observed in pre-vaccination and sham controls (*P*<0.01). Frequency of intracellular TNF-α expressing CD4^+^ T cell response was also found significantly different between vaccinated and sham monkeys (*P*<0.05; Fig. S7, A and B), but intracellular TNFα^+^ CD8^+^ or IL-2^+^ CD4^+^/CD8^+^ T cell response to S peptides was not statistically different between groups (*P*>0.05; Fig. S7, C-F).

**Fig. 6.**
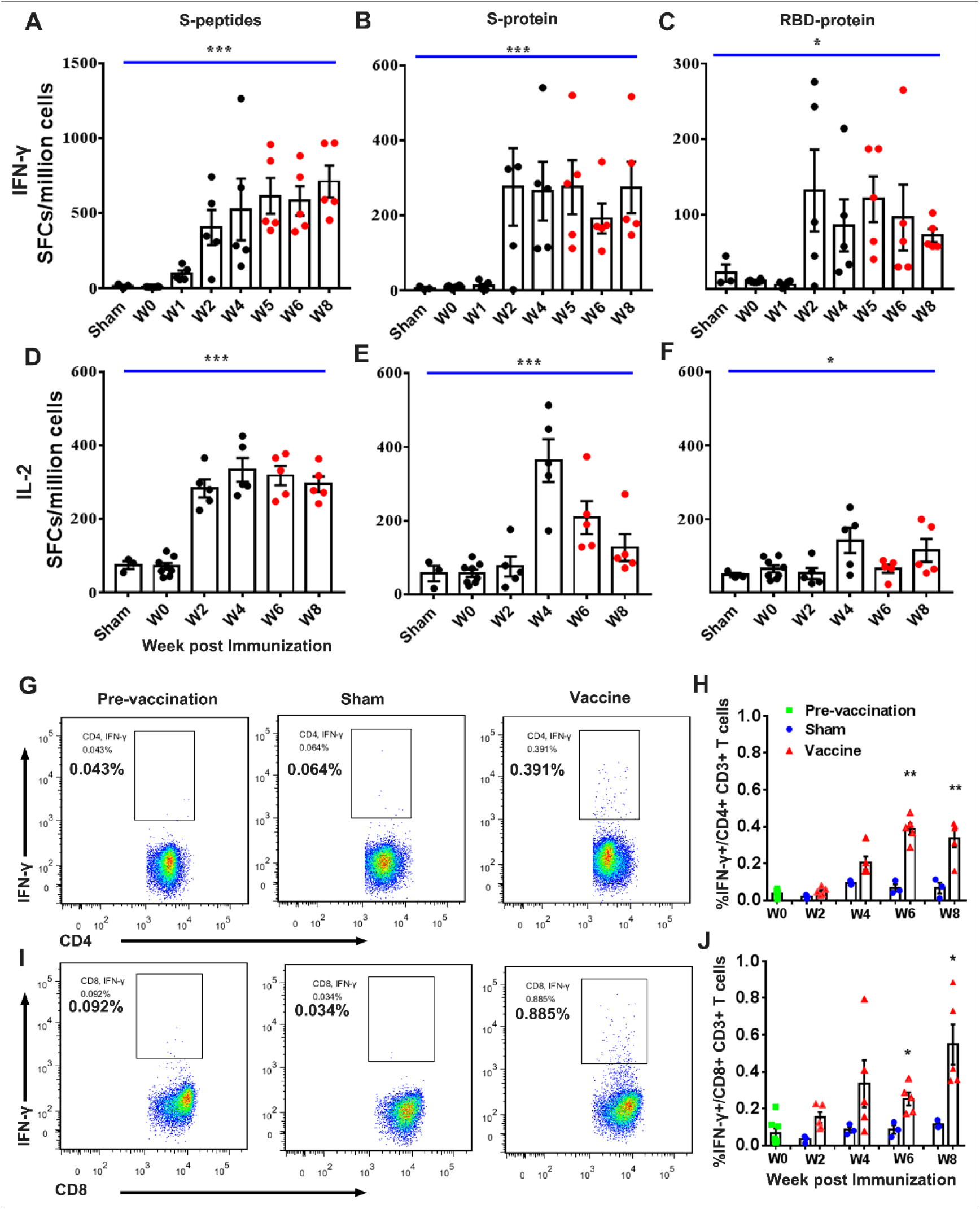
Specific T cell response of PBMCs from Rhesus macaques immunized with Sad23L-nCoV-S and Ad49L-nCoV-S vaccines or sham controls by prime-boost vaccination regimen. (**A-C**) IFN-γ or (**D-F**) IL-2 secreting T cell response (SFCs/million cells) to S peptides, S or RBD protein was measured by ELISpot. (**G-J**) Frequency of intracellular IFN-γ expressing CD4+/CD8+ T cell responses to S peptides was determined by ICS, respectively. Data are shown as mean ± SEM. P values are calculated with two-tailed t test. Statistically significant differences are shown with asterisks (*, *P*<0.05; **, *P*< 0.01 and ***, *P*< 0.001).

In summary, prime-boost vaccination with Sad23L-nCoV-S and Ad49L-nCoV-S vaccines at an interval of 4 weeks elicited higher levels of specific antibody and T-cell responses against SARS-CoV-2 in rhesus macaques, which was recommended as COVID-19 vaccine candidates for clinical trials in humans.

### Biodistribution of Sad23L-nCoV-S and Ad49L-nCoV-S vaccine strains in the tissues of inoculated mice

Neutralizing antibody titers to Sad23L and Ad49L vectors were determined in rhesus macaques and mice after prime only and prime-boost vaccinations (Fig. 7, A and B; Fig. S8). A high NAb reactivity to an individual adenoviral vector of either Sad23L or Ad49L in macaques (1:1280 or 1:640) and mice (1:576 or 1:352) was induced post prime immunization, but not enhanced post boost vaccination with a heterologous vector (Fig. 7, A and B), which might limit the homologous boosting of adenovirus vectored vaccines. The lung, spleen, liver and muscle tissues (at intramuscular injection site and para-tissues) from prime only or prime-boost immunized C57BL/6 mice were examined 4 weeks after inoculation of vaccines in order to assess vaccine delivery by amplifying specific adenoviral hexon sequences of both Sad23L and Ad49L vectors. The predicted 500bp PCR band was detected in lung, spleen and liver tissues after Sad23L-nCoV-S vaccination, while the PCR band was observed in lung and liver tissues but not found in the spleen after Ad49L-nCoV-S vaccination (Fig. 7C). The expression of S antigen was observed in splenocytes and hepatocytes from prime-only Sad23L-nCoV-S or Ad49L-nCoV-S and prime-boost vaccines inoculated C57BL/6 mice by immunofluorescence staining (Fig. 7D), but not found in lung and muscle tissues (Fig. S9).

**Fig 7.**
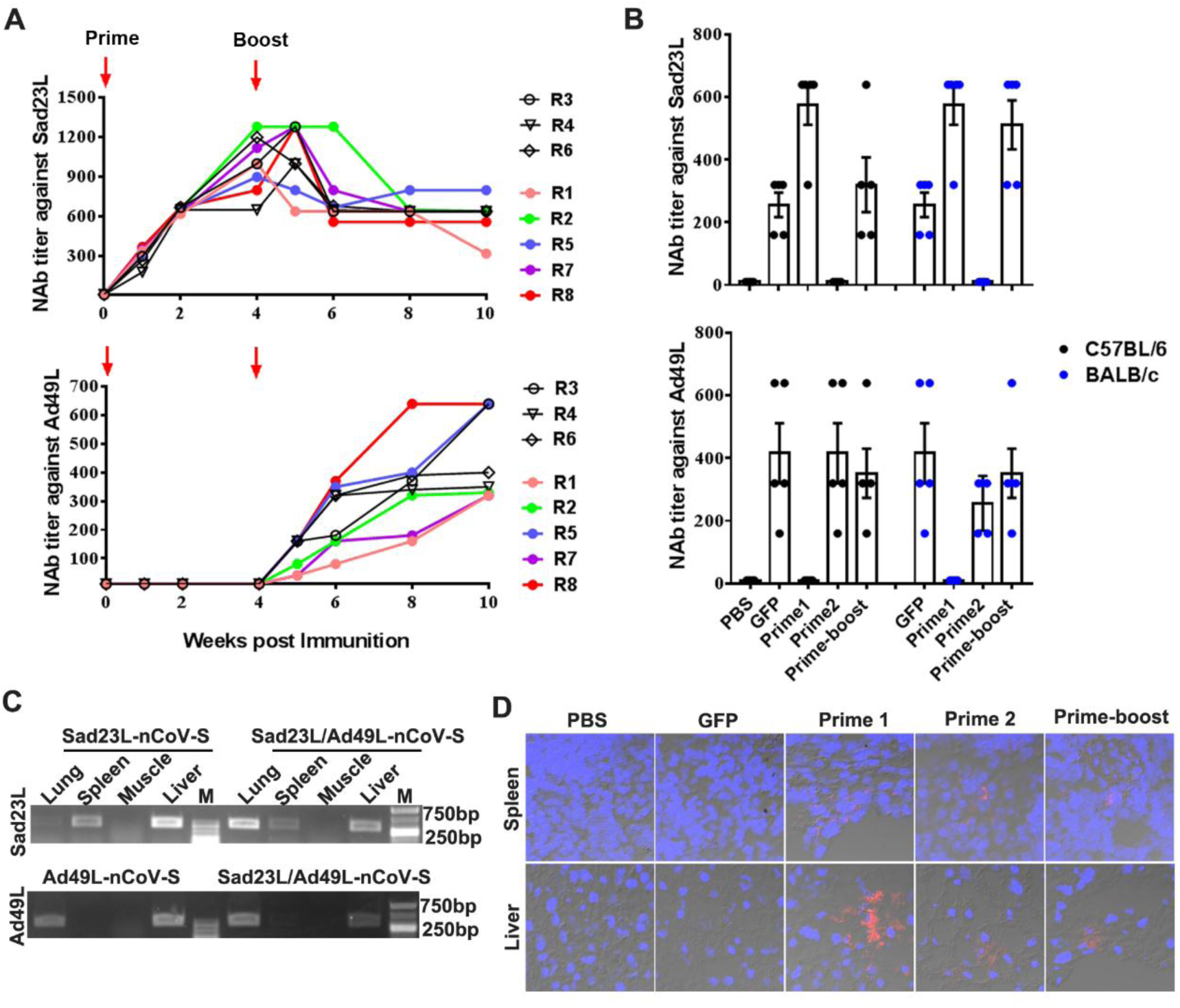
Biodistribution of Sad23L-nCoV-S and Ad49L-nCoV-S vaccines in inoculated animals. (**A**) Serum NAb titers to Sad23L and Ad49L vectors were measured in macaques immunized by prime-boost inoculation with two vaccines at 4 week interval, or (**B**) in C57BL/6 and BALB/c mice 4 weeks post prime only or prime-boost vaccination with two vaccines or vectorial controls. (**C**) Nested-PCR amplification of Sad23L or Ad49L-hexon gene (500bp) in tissues of C57BL/6 mice 4 weeks after inoculation by prime only or prime-boost immunization with Sad23L-nCoV-S and Ad49L-nCoV-S vaccines. (**D**) Expression of S protein in splenocytes and hepatocytes of tissue frozen sections from vaccine immunized or control C57BL/6 mice by immunofluorescence staining with a human monoclonal antibody to SARS-CoV-2 S and DAPI.

## DISCUSSION

A safe and effective COVID-19 vaccine is one of the most wanted goods in the world. Among 33 COVID-19 candidate vaccines in clinical trials (WHO report on 28 August 2020) (*6*), 9 vaccines have initiated clinical phase III trial, and experimental data from pre-clinical and/or clinical studies have been published (*8–21*). In the frontline of COVID-19 vaccine candidates in clinical trial phase III, the range of approaches includes three inactivated virus, two lipid nanoparticle (LNP)-encapsulated mRNA and four non-replicating adenovirus vectored vaccines (*6*). Beyond general comments on the disadvantages of major types of COVID-19 vaccines (*24, 25*), the limitations of inactivated SARS-CoV-2 vaccines are crucial biosafety concerns regarding large amounts of infectious virus being processed above biological safety level-3 (BSL-3) condition and production capacity; for LNP-mRNA vaccines instability and lack of evidence for efficacy are puzzling; for adenovirus vectored vaccines, pre-existing immunity to the carrier virus and the relative weakness of single-shot immunization appear limiting factors.

In this study, we generated two novel adenovirus vectored COVID-19 vaccines encoding the full-length S gene of SARS-CoV-2. The intact S glycoprotein rather than the shorter S or RBD proteins was shown to be the most effective antigen eliciting protective immunity against SARS-CoV-2 infection in DNA vaccines and Ad26 vectored vaccines (*12,13,26*). The vectors Sad23L and Ad49L originated from simian adenovirus type 23 and human adenovirus type 49, respectively (*22, 23*), are novel adenoviral vector. Comparing Ad5-vectored COVID-19, ChAdox1 nCoV-19 and Ad26.COV2.S vaccines (*8-12,14,21*), three attractive aspects emphasized in this study are highlighted below.

Firstly, there is a low-seroprevalence of pre-existing antibodies to Sad23L and Ad49L vectors in humans. According to the investigation of NAb to three types of adenoviruses in Chinese population, the prevalence of both Sad23L and Ad49L NAb was below 10%, while the prevalence of Ad5 NAb was over 75% (Fig. 1D). However, the pre-existing anti-Ad5 immunity might partly limit vaccine effectiveness, especially for populations aged over 50 (*8,9,24,25*), while the low seroprevalence of antibodies to novel adenoviral vectors such as ChAdox1 (*11*), Ad26 (*12,14,21,27*), and Sad23L and Ad49L used in this study might avoid a negative impact on vaccine efficacy (*22–24*).

Secondly, the prime-boost vaccination regimen with two heterologous adenoviruses vectored Sad23L-nCoV-S and Ad49L-nCoV-S vaccines examined in mice and rhesus macaques suggested that the boost with Ad49L-nCoV-S vaccine significantly enhanced the levels of neutralizing antibody and IFN-γ expressing CD4^+^/CD8^+^ T cell responses following prime-immunization with Sad23L-nCoV-S vaccine (Fig. 4-6). In addition, specific IFN-γ secretion T cell response was prolonged at high level after boosting vaccination. Two recent publications compared prime only with a single-dose of ChAdOx1 nCoV-19 and homologous boost with a second dose of ChAdOx1 nCoV-19 vaccine in mice and pigs, and human clinical trials, showed that boost vaccination significantly increased and prolonged the level of binding or neutralizing antibody response to SARS-CoV-2 (*11, 20*). By boosting with Ad5-S vaccine the Ad26-S vaccine primed immunization, a higher level of immunity was observed in humans as reported in a recent Russian study (*21*). Compared with homologous boosting of ChAdOx1 nCoV-19 or boosting of Ad5-S vaccines (*11, 21*), heterologous prime-boost vaccinations with Sad23L-nCoV-S and Ad49L-nCoV-S vaccines have the advantage of avoiding vector’s immunity interfering with prime vaccination or enhancing pre-existing Ad5 immunity (*22–25*).

Thirdly, in order to achieve an effective vaccine immunity, a low dose of two heterologous adenovirus vectored vaccines (<5×10^10^ vp) with prime-boost immunization regimen should theoretically reduce severe adverse reaction induced by a high dose of adenovirus vectored vaccine (>5×10^10^ vp) with prime only immunization in clinical trials (*8,9,11,14*). In this study, a relatively low dose of Sad23L-nCoV-S and Ad49L-nCoV-S vaccines elicited a robust immunity in both young mice and older rhesus macaques (aged 11-14 years), but no obvious clinical symptoms or histopathological changes were observed (Fig. 2; Fig. S1; Fig. S2).

There are two limitations to this work. Considering biosafety and restriction for SARS-CoV-2 virus manipulation, conventional live virus neutralization test (cVNT) and virus injection challenge experiments have not been performed in this study. However, sVNT and pVNT reported in this study have well demonstrated a significant correlation with cVNT in NAb titers to SARS-CoV-2 (*P*<0.0001, *R*=0.7678-0.8591) (*28*). This was also evidenced by close correlation between ELISA and pVNT or cVNT by other studies using adenovirus vectored COVID-19 vaccines in rhesus macaques (*P*<0.0001, *R*=0.8314-0.8427) (*12*), hamsters (*P*<0.0001; *R*=0.7849) (*13*) and human clinical phase II trial (*P*<0.0001, *R*=0.72-0.75) (*9*). In addition, compared with a panel of convalescent serum samples from COVID-19 patients the level of binding or neutralizing antibodies observed in this study was favorable and in line with all other published data from COVID-19 vaccines. Protective efficacy of Sad23L-nCoV-S and Ad49L-nCoV-S vaccines for prime-boost vaccination of animals might be referred to ChAdOx1 nCoV-19 or Ad26.COV2.S vaccine immunized rhesus macaques against SARS-CoV-2 challenge (*10, 12*), because Sad23L and ChAdOx1, Ad49L and Ad26 were classified in the same species E or D of adenoviruses, respectively (*7*).

In conclusion, two novel adenovirus vectored COVID-19 vaccines were produced, which could, when used in succession, safely elicit robust humoral and T-cell immune response to SARS-CoV-2 in mice and Rhesus macaques. Prime-boost vaccination regimen with priming of Sad23L-nCoV-S and boosting of Ad49L-nCoV-S vaccines were recommended to develop the better protective immunity against SARS-CoV-2 in humans. These two vaccines are being planned for clinical phase I/II trials after an extensive safety evaluation has been carried out in pre-clinical animal examination.

## MATERIALS AND METHODS

### Rhesus macaques and ethics statement

Eight healthy outbred male rhesus macaques (*Macaca mulatta*) aged 11-14 years were randomly allocated to this study (Table S1). Experimentation and sample collection were ethically approved and carried out by the Huazheng Laboratory Animal Breeding Centre, Guangzhou, China. All animal care and experimental procedures (NFYYLASOP-037) were in accordance with national and institutional policies for animal health and wellbeing.

The welfare issues (housing, feeding, environmental enrichment, etc.) were in accordance with the recommendations of the Weatherall report (https://acmedsci.ac.uk/more/news/the-use-of-non-human-primates-in-research). Animals were individually housed in spacious cages and were provided with commercial food pellets supplemented with appropriate treats. Drinking water was provided ad libitum from an automatic watering system. Animals were monitored daily for health and discomfort. Blood samples were obtained using sterilized needle and syringe from the venous vessels of animal legs.

### Cells and mice

HEK-293A, HEK-293T and HEK293T-hACE2 cells (Sino Biological) were maintained in complete Dulbecco’s modified Eagle’s medium (DMEM, Gibco) supplemented with 10% fetal bovine serum (FBS, Gbico) and incubated at 37°C in 5% CO_2_.

Female C57BL/6 and BALB/c mice were obtained from the Animal Experimental Centre of Southern Medical University, Guangdong, China.

### COVID-19 patients’ serum samples

A total of 48 convalescent serum samples were provided by The First People’s Hospital of Foshan, Shenzhen or Guangzhou Center for Disease Control and Prevention (CDC), China. The samples were collected from 25 asymptomatic, 14 mild and 9 severe COVID-19 infected subjects. All serum samples were inactivated for 40 min by heating at 56°C in the water bath.

### Construction of two novel adenovirus vectored COVID-19 vaccine strains

According to the description of Sad23L vector (SAdV23, GenBank: AY530877.1) (*22, 23*), the replication defective adenoviral vector Ad49L was constructed by deleting the E1 and E3 regions of the full-length human adenovirus serotype 49 genome (HAdV49, GenBank: DQ393829.1). The E4 region open reading frame 6 (orf6) was replaced by the corresponding element of human adenovirus type 5 (Ad5) in order to improve virus propagating efficiency. The full-length S protein gene of SARS-CoV-2 (GenBank: MN908947.3) was optimized and synthesized (Beijing Genomics Institute, China) and was cloned into adenoviral vectors Sad23L and Ad49L, respectively. The recombinant adenoviral constructs Sad23L-nCoV-S and Ad49L-nCoV-S were rescued, and the novel adenovirus vectored COVID-19 vaccine strains were propagated from HEK-293A packaging cells. The vaccine strains were serially passaged for stability by 12 generations when the full cytopathic effect appeared. Virus purification was performed by cesium chloride density gradient centrifugation as previously described (*23*).

### Histopathological examination

Mice tissues were stained with hematoxylin and eosin (H&E) and examined microscopically for histopathological changes by Guangzhou Huayin Medical Science Company Limited (Guangzhou, China).

### Western blotting

HEK-293A cells were infected with Sad23L-nCoV-S and Ad49L-nCoV-S strains, respectively, and Sad23L-GFP and Ad49L-GFP vectorial viruses were used as mock control. The expression of SARS-CoV-2 S protein was analyzed by Western blotting with rabbit polyclonal antibody to SARS-CoV-2 RBD (Sino Biological, China) and heat-inactivated human serum samples from Chinese COVID-19 infected patients. Glyceraldehyde-3-phosphate dehydrogenase (GADPH) was included as a loading control. The membranes were washed five times and developed by Supersignal West Pico Plus chemiluminescent substrate (Thermo Scientific, USA).

### Immunofluorescence staining

Cells infected with vaccine strains or vector control viruses were fixed in cell culture plates, while tissues were collected from vaccinated and control mice. Cell layers or tissue frozen sections were incubated with human monoclonal antibody to SARS-CoV-2 RBD (OkayBio, China), and then washed with PBST. Anti-human IgG-Alexa Fluor 594 antibody (Thermo Scientific, USA) in 1% BSA-PBST was added to the cells for 30 min at 37°C. DAPI was added to stain cell nuclei.

### Adenovirus neutralizing antibody assay

Human plasma samples were collected from 600 healthy blood donors at six blood centers (100 per center) across China, including Shenzhen (south), Guangzhou (south), Yichang (central), Harbin (northeast), Chengdu (southwest) and Xian (west) blood centers. Plasmas were tested on HEK-293A cells for neutralizing Sad23L-GFP, or Ad49L-GFP vectorial viruses by green fluorescent activity assay as previously described (*22, 29*).

### Animal immunization

Female C57BL/6 mice (5-6 weeks, n=5 each group) were individually inoculated intramuscularly (i.m.) with a dose of 10^7^, 10^8^ and 10^9^ PFU Sad23L-nCoV-S or Ad49L-nCoV-S vaccine, respectively. A dose of 10^9^ PFU Sad23L-GFP or Ad49L-GFP vectorial virus and an equivalent volume of PBS were used as sham control.

Female C57BL/6 and BALB/c mice (5-6 weeks, n=5 each group) were prime inoculated intramuscularly with a dose of 10^9^ PFU Sad23L-nCoV-S vaccine, and then at 4 week interval were boosted with a dose of 10^9^ PFU Ad49L-nCoV-S vaccine. A dose of 10^9^ PFU Sad23L-GFP and a dose of 10^9^ PFU Ad49L-GFP were used as sham control.

Five rhesus macaques aged 11 to 14 years (Table S1) were first injected intramuscularly with a dose of 5×10^9^ PFU Sad23L-nCoV-S vaccine, and then at 4 week interval received a second dose of 5×10^9^ PFU Ad49L-nCoV-S vaccine. Three rhesus macaques aged 11 to 13 years (Table S1) were vaccinated by prime-boost regimen with a dose of 5×10^9^ PFU Sad23L-GFP and a dose of Ad49L-GFP vectorial adenoviruses as sham control.

### Enzyme-linked immunosorbent assay (ELISA)

The microtiter plates (Corning, USA) were coated overnight with 2μg/ml of SARS-CoV S or RBD proteins (Sino Biological, China). Serum samples were 2-fold serially diluted and S or RBD binding antibody (S-BAb or RBD-BAb) were detected by ELISA. Secondary antibodies were goat anti-mouse IgG-HRP (Beijing Bersee Science and Technology, Co. LTd, China), rabbit anti-monkey IgG-HRP (Bioss, China) and goat anti-human IgG-HRP conjugates (Sigma, USA), respectively. Endpoint titers were defined as the highest reciprocal serum dilution that yielded an absorbance > 0.2 and a ratio of signal than cutoff (S/CO) > 1. Log10 end point titers were reported (*26*).

Goat anti-mouse IgG1, IgG2a, IgG2b or IgG3 heavy chain-HRP conjugates were used for IgG sub-classification according to manufacturer’s instruction (Abcam, UK).

### Surrogate virus neutralization test (sVNT)

The surrogate virus based neutralization test (sVNT) was used for measuring neutralizing antibody (NAb) to SARS-CoV-2 as previously described (*28*). Briefly, microtiter plates (Corning) were coated overnight with 2μg/ml hACE2 protein (Sino Biological, Beijing, China) at 4 °C, followed by blocking with OptEIA assay diluent (BD). The HRP–RBD conjugate (3 ng) was pre-incubated with 100 μl of diluted serum samples for 1 h at 37 °C, then added into hACE2 pre-coated plate for 1 h at room temperature. Plates were washed five times by PBST. A colorimetric signal was developed with TMB, and equal volume of stop solution was added to terminate the reaction. The absorbance reading was performed at 450 nm and 570 nm. Inhibition rate (%) = (1 − sample optical density value/negative control optical density value) × 100.

### Pseudovirus neutralization test (pVNT)

Pseudoviruses expressing a luciferase reporter gene were generated for measuring of NAb to SARS-CoV-2 as previously described (*26*). Briefly, the packaging construct psPAX2 (Addgene), plasmid pLenti-CMV Puro-Luc (Addgene), and pcDNA3.1-SARS-CoV-2 SΔCT (deletion of the cytoplasmic tail) were co-transfected into HEK-293T cells. The supernatants were collected 48 hours post-transfection and the pseudoviruses were purified by filtration through 0.45 µm filter. The pVNT titers were measured with HEK293T-hACE2 cells in 96-well tissue culture plates. Two-fold serial dilutions of heat-inactivated serum samples were prepared and mixed with 50 µl of pseudovirus. After incubation at 37°C for 1h, the serum-virus mixture was added to HEK293T-hACE2 cells. Forty-eight hours after infection, cells were lysed in Steady-Glo Luciferase Assay (Promega) according to the manufacturer’s instructions. The pVNT titer of SARS-CoV-2 antibody was defined as the sample dilution at which a 50% inhibition rate. Inhibition rate (%) = (1 − sample RLU/virus control RLU) × 100. Log10 pVNT(ID_50_) titer was reported.

### ELISpot

Monkey (or mouse) IFN-gamma, IL-2 or IL-4 ELISpotPLUS kits (MabTech) were used to determine SARS-CoV-2 S antigen-specific T lymphocyte response. Rhesus macaque’s PBMCs (2×10^5^ cells/well) or mouse splenocytes (5×10^5^ cells/well) were stimulated with S peptides, S or RBD protein (5μg/ml) in triplicates, respectively. Seventy-nine peptides encoding for amino acid sequence of SARS-CoV-2 S protein were predicted (http://www.iedb.org/) and synthesized by Guangzhou IGE Biotechnology LTD (Table. S3). Spots were counted with a CTL Immunospot Reader (Cellular Technology Ltd). The results were expressed as spot forming cells (SFCs) per million cells.

### Intracellular cytokine staining (ICS) and flow cytometry

Mouse splenocytes (2×10^6^ cells/well) or monkey PBMCs (1×10^6^ cells/well) were stimulated with S peptide pools (3μg/ml each peptide), or medium as negative control, in triplicates. After 4h, the cells were incubated with Golgi Plug (BD) for 12h at 37°C. Cells were collected and stained with anti-mouse or anti-monkey CD3, CD4 and CD8 surface marker antibodies (BD). Cells were fixed with IC fixation buffer, permeabilized with permeabilization buffer (BD) and stained with anti-mouse or anti-monkey interferon-γ (IFN-γ), interleukin-2 (IL-2) and tumor necrosis factor α (TNF-α) (BD). All samples were tested with BD FACS Canton flow cytometer (BD).

### Nested-PCR amplification

Genomic DNA was extracted from homogenized tissue of adenovirus vectored vaccine inoculated mice by High Pure Viral Nucleic Acid Kit (Roche). DNA fragments of Sad23L and Ad49L vectors were amplified by nested PCR with primers specific to adenoviral hexon sequences (Table S4) (*23*).

### Statistical analyses

Data are analyzed with unpaired two-tailed *t* test, one-way ANOVA. Neutralizing antibody titer data were log transformed before analysis. Neutralizing antibody titer data generated by the RBD-BAb and sVNT or S-BAb and pVNT assays were compared using Spearman nonparametric correlation. Statistically significant differences are indicated with asterisks (* *P* <0.05; ** *P* <0.01 and *** *P* <0.001). All graphs are generated with GraphPad Prism 7 software.

## ACKNOWLEDGMENTS

The authors thank professors Kwok-Yung Yuen from The University of Hong Kong for his assistance for purchasing simian adenoviruses type 23 and human adenovirus type 49 strains; Dongming Zhou from The Institut Pasteur in Shanghai for his technical help for construction of adenovirus vectors; Shuwen Liu and Mengfeng Li from Southern Medical University for the help for pseudovirus neutralization test, study design and writing of manuscript; Caiping Guo from Shenzhen Weiguang Biological Products Co., Ltd for her helpful suggestion on identification of vaccine strains. The authors also thank The First People’s Hospital of Foshan, Shenzhen and Guangzhou CDC for providing COVID-19 patients’ serum samples; Shenzhen, Guangzhou, Yichang, Harbin, Chengdu and Xian Blood Centers for providing plasma samples of blood donors.

## Author contributions

C.L., L.S., T.L. and Y.Z. designed the research. L.S., P.Z., B.L., C.Y., C.L.L., Q.W., L.Z. and T.L. carried out the experiments. X.T., J.L., S.H., J.Z. and Y.F. provided COVID-19 patient or healthy blood donor samples. S.L., C.L., Y.Z. and J.P.A. analyzed data and wrote the paper.

## Data and materials availability

All data associated with this paper can be found in the main text or the supplementary materials. Two novel adenoviral vectors (Sad23L and Ad49L) are available under a material transfer agreement with BRK upon request to the corresponding author.

## Supplementary Materials

**Fig. S1.**
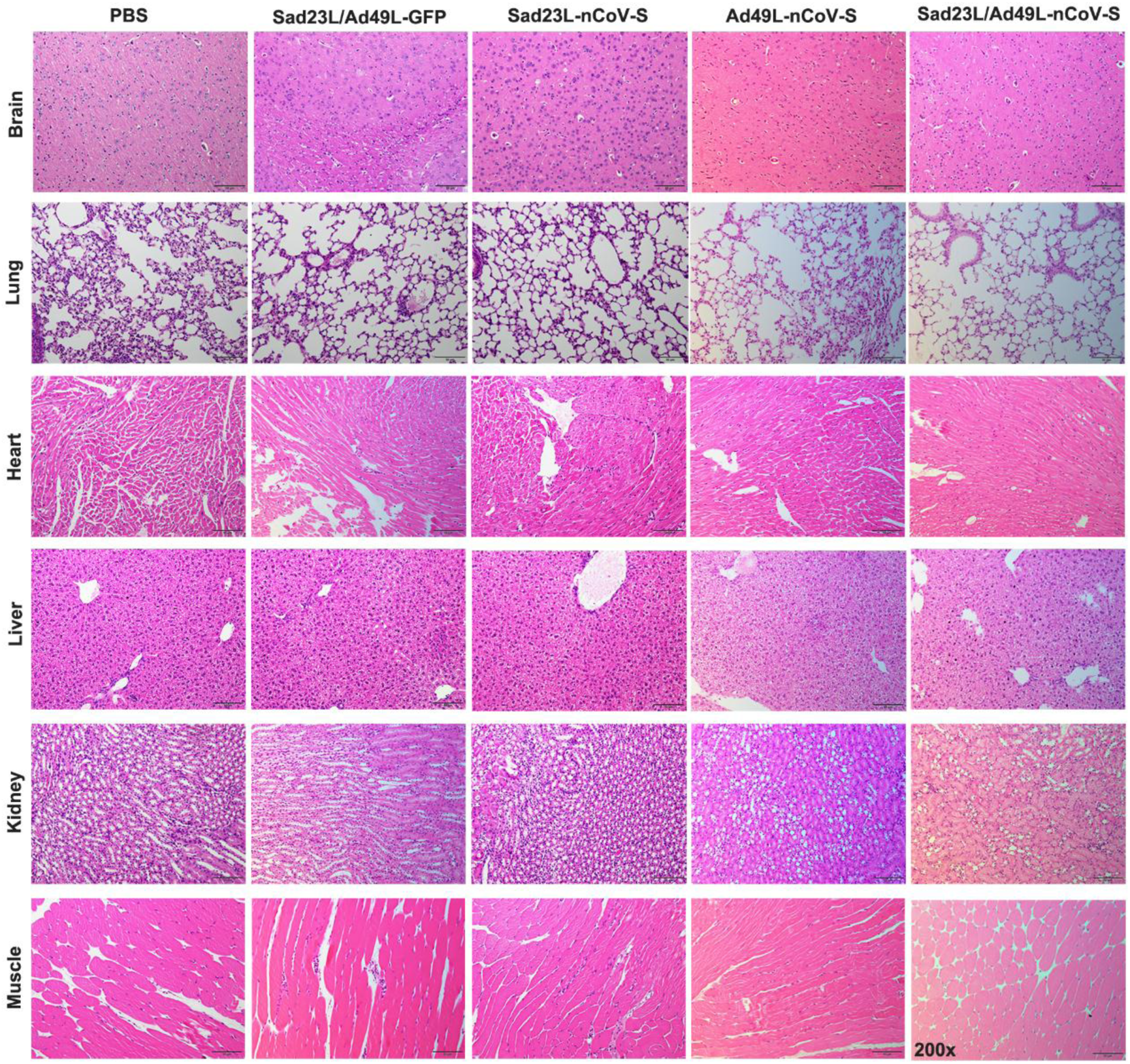
Histopathological examination from Sad23L-nCoV-S and Ad49L-nCoV-S vaccines inoculated C57BL/6 mice. Histopathological examination was carried out for brain, lung, heart, liver, kidney and muscle tissues (at intramuscular injection site and para-tissues) of mice in 4 weeks post prime only or prime-boost immunization with these two vaccines. Tissues were stained with hematoxylin and eosin.

**Fig. S2.**
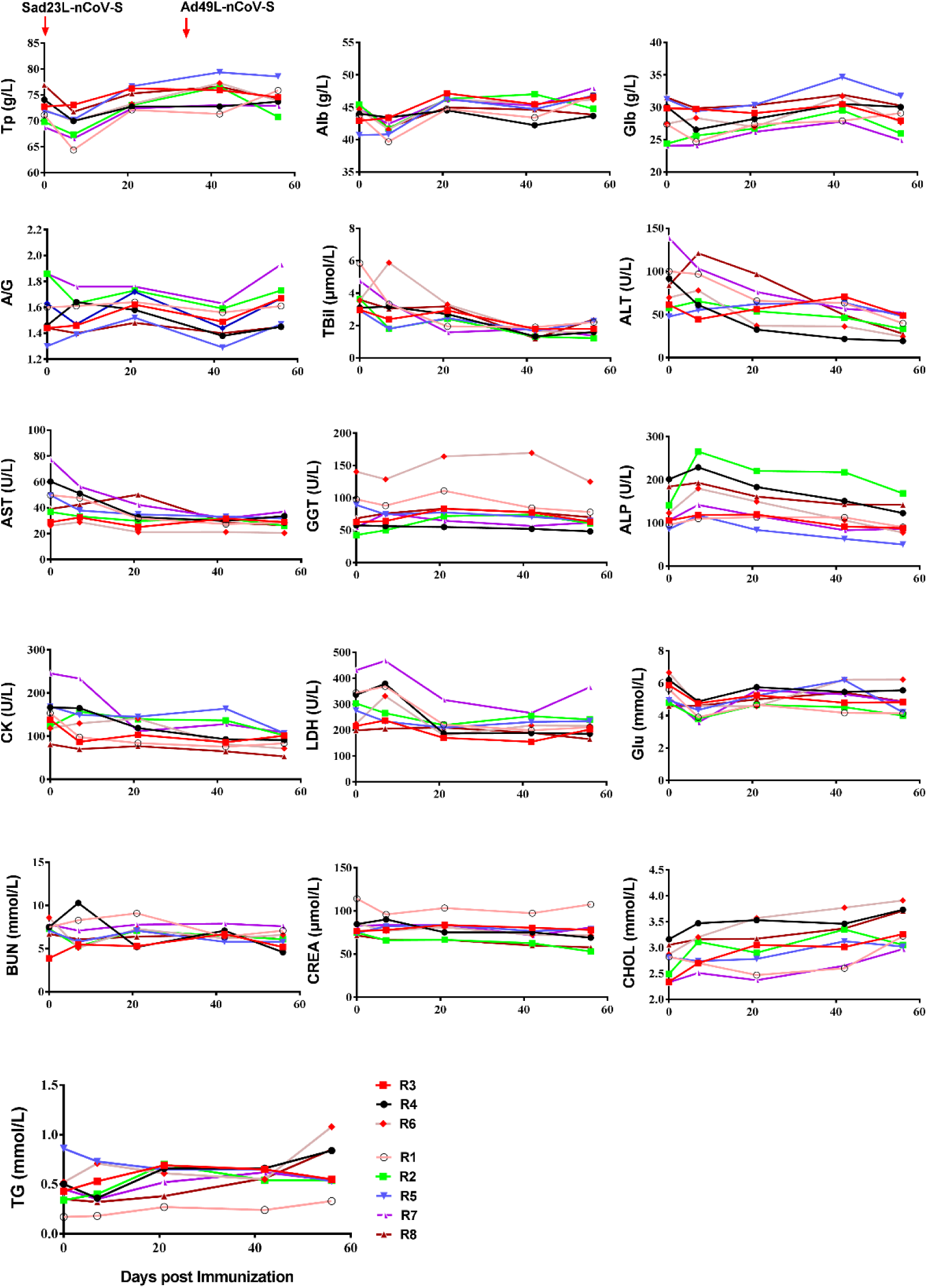
Kinetic change of hematological and clinical biochemistry indexes during the course of vaccinated or sham control rhesus macaques. Rhesus macaques were intramuscularly inoculated by prime-boost immunization with Sad23L-nCoV-S and Ad49L-nCoV-S vaccines at an interval of 4 weeks. A panel of hematological and biochemical indexes were measured from blood samples at different time points. TP, Total protein; Alb, Albumin; Glb, Globulin; A/G, Albumin/globulin ratio; TBil, Total bilirubin; ALT, Alanine aminotransferase; AST, Aspartate aminotransferase; GGT, γ–glutamyltranspeptidase; ALP, Alkaline phosphatase; CK, Creatine kinase; LDH, Lactate dehydrogenase; Glu, Glucose; BUN, Blood urea nitrogen; CREA, Creatinine; CHOL, Total cholesterol; TG, Triglycerides. R1-R8 indicate rhesus macaques.

**Fig. S3.**
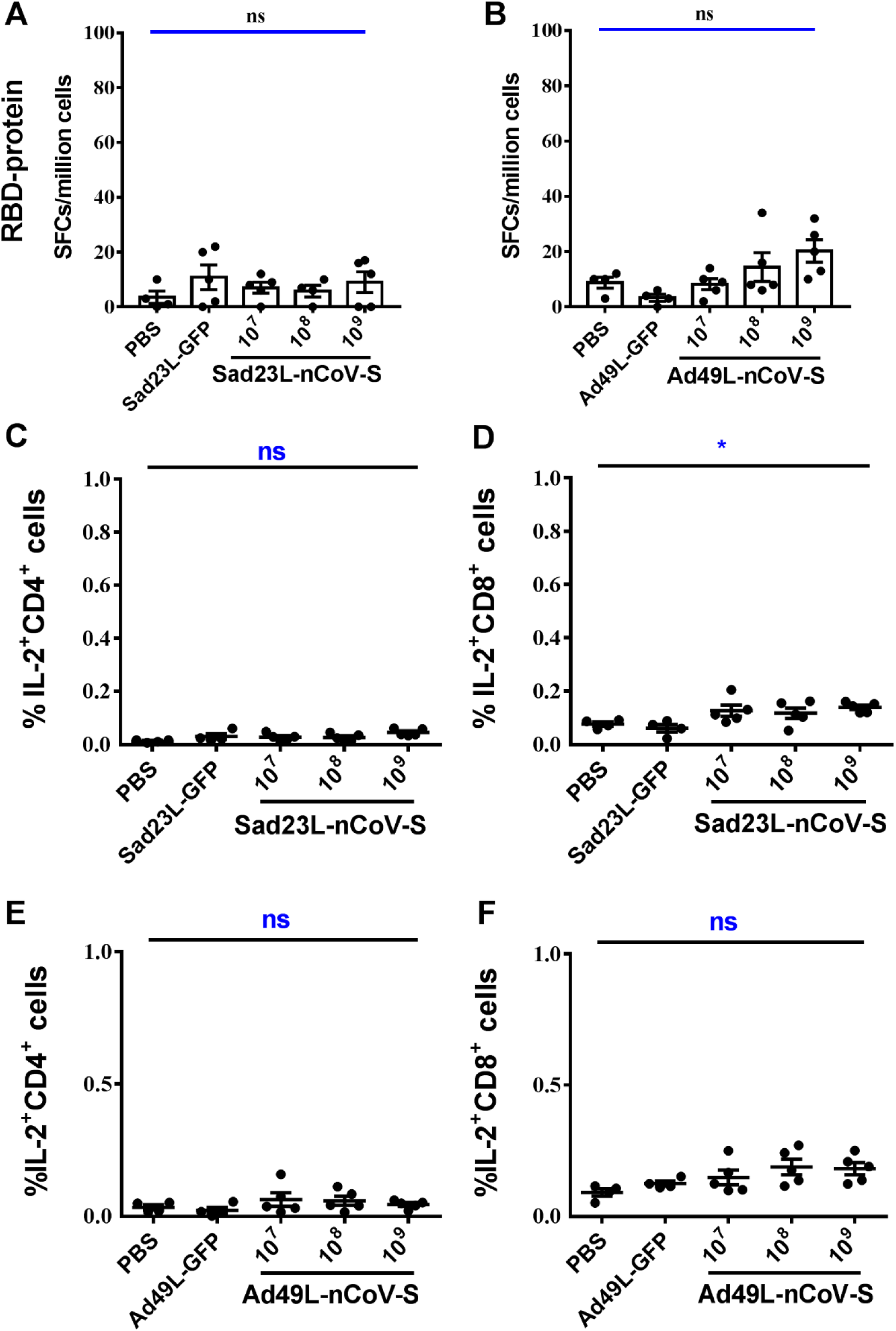
Specific T cell response of splenocytes from C57BL/6 mice (n=5/group) immunized with a single dose of Sad23L-nCoV-S or Ad49L-nCoV-S vaccine. (**A** and **B**) Specific IFN-γ secreting T cell response (spot forming cells [SFCs] per million cells) to RBD protein was measured by ELISpot, respectively. (**C** and **D**) Frequency of IL-2^+^ CD4^+^ or CD8^+^ T cell response to S peptides from mice elicited by Sad23L-nCoV-S vaccines was measured by ICS, respectively. (**E** and **F**) Frequency of IL-2^+^ CD4^+^ or CD8^+^ T cell response to S peptides from mice immunized by Ad49L-nCoV-S vaccines was measured by ICS, respectively. Data were shown as means ± SEM (standard errors of means). *P* values were analyzed by one-way ANOVA with 2-fold Bonferroni’s test. Statically significant differences were showed with asterisks (*, *P*<0.05; **, *P*< 0.01 and ***, *P*< 0.001). ns, *P*>0.05 and no significant difference.

**Fig. S4.**
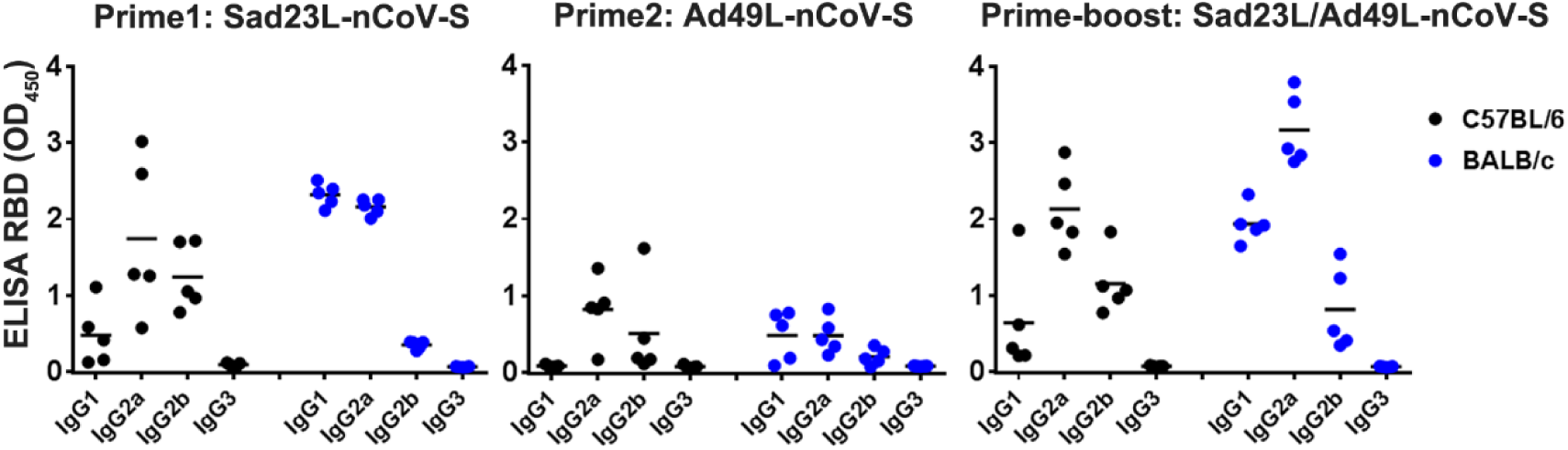
IgG subclass antibodies against RBD protein in sera of C57BL/6 and BALB/c mice immunized by prime only or prime-boost with Sad23L-nCoV-S and Ad49L-nCoV-S vaccines. The sera were collected 4 weeks post prime only or prime-boost immunized C57BL/6 and BALB/c mice. IgG subclass antibodies in sera were detected against RBD protein by ELISA with secondary antibody-HRP conjugate specific to mouse IgG subclass.

**Fig. S5.**
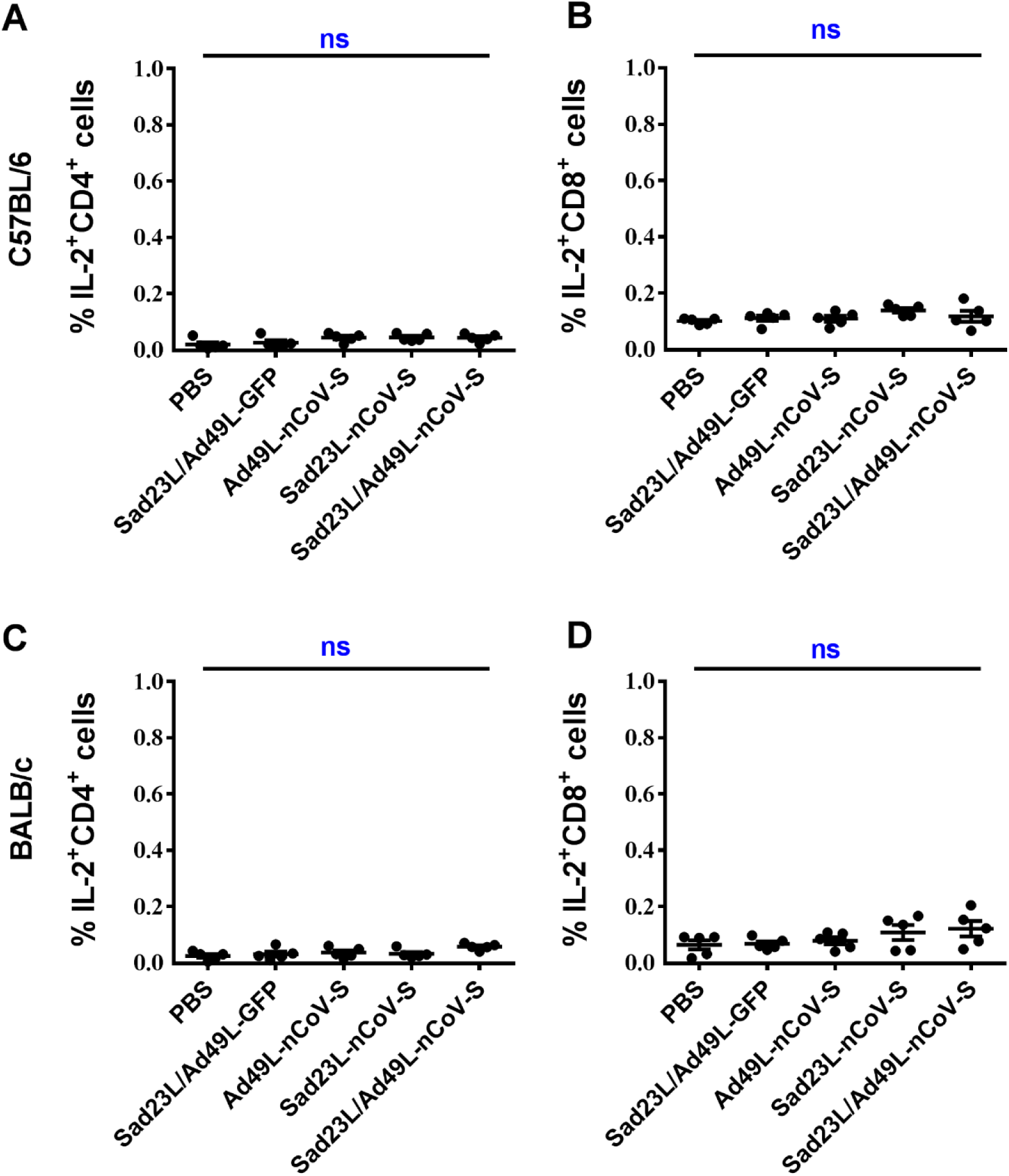
Frequency of IL-2 expressing CD4+/CD8+ T cell responses of splenocytes from prime-boost immunized C57BL/6 and BALB/c mice with Sad23L-nCoV-S and Ad49L-nCoV-S vaccines. (**A** and **B**) Frequency of IL-2^+^ CD4^+^ or CD8^+^ T cell responses of splenocytes to S peptides from C57BL/6, or (**C** and **D**) from BALB/c mice. Data were shown as means ± SEM (standard errors of means). *P* values were analyzed by one-way ANOVA. ns, *P*>0.05 and no significant difference.

**Fig. S6.**
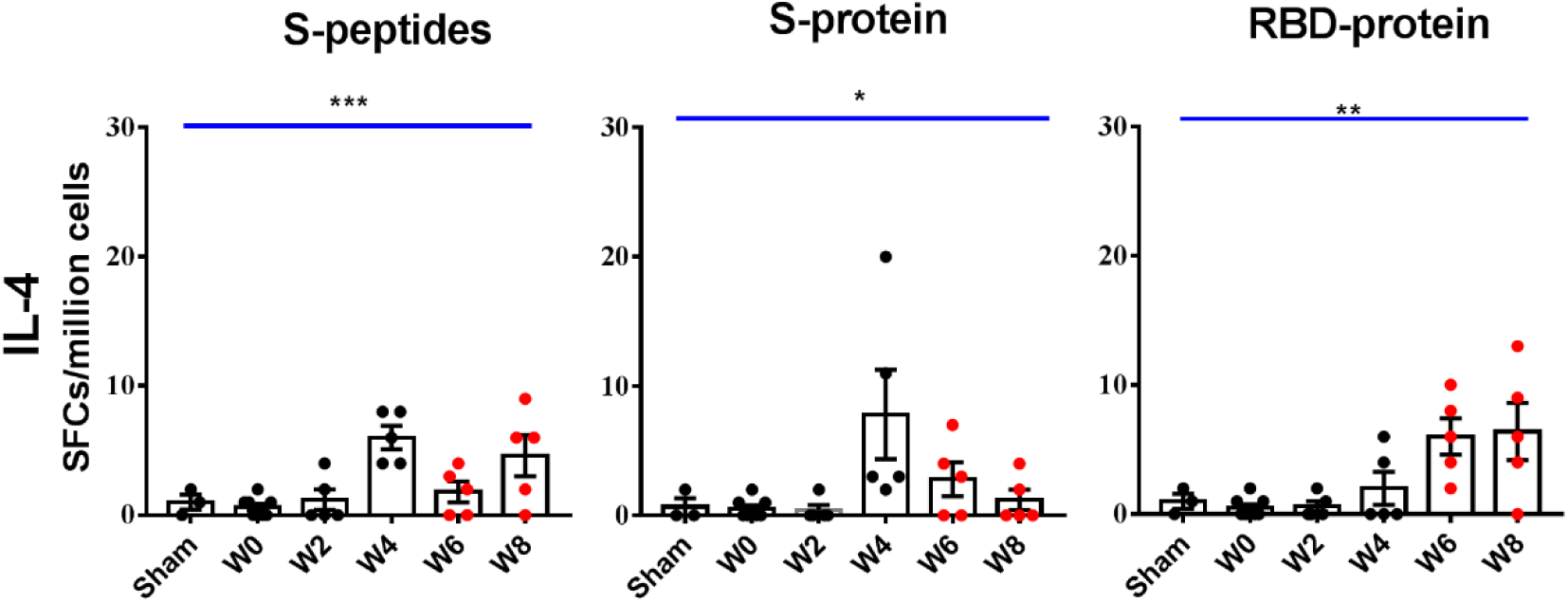
IL-4 secreting T cell response in prime-boost immunized rhesus macaques with Sad23L-nCoV-S and Ad49L-nCoV-S vaccines. The number of specific IL-4 secreting T cells (SFCs/million cells) to S peptides, S or RBD protein in PBMCs of monkeys was measured by ELISpot, respectively. Data were shown as means ± SEM (standard errors of means). *P* values were analyzed by one-way ANOVA with 2-fold Bonferroni’s test. Statically significant differences were showed with asterisks (*, *P*<0.05; **, *P*< 0.01 and ***, *P*< 0.001). ns, *P*>0.05 and no significant difference.

**Fig. S7.**
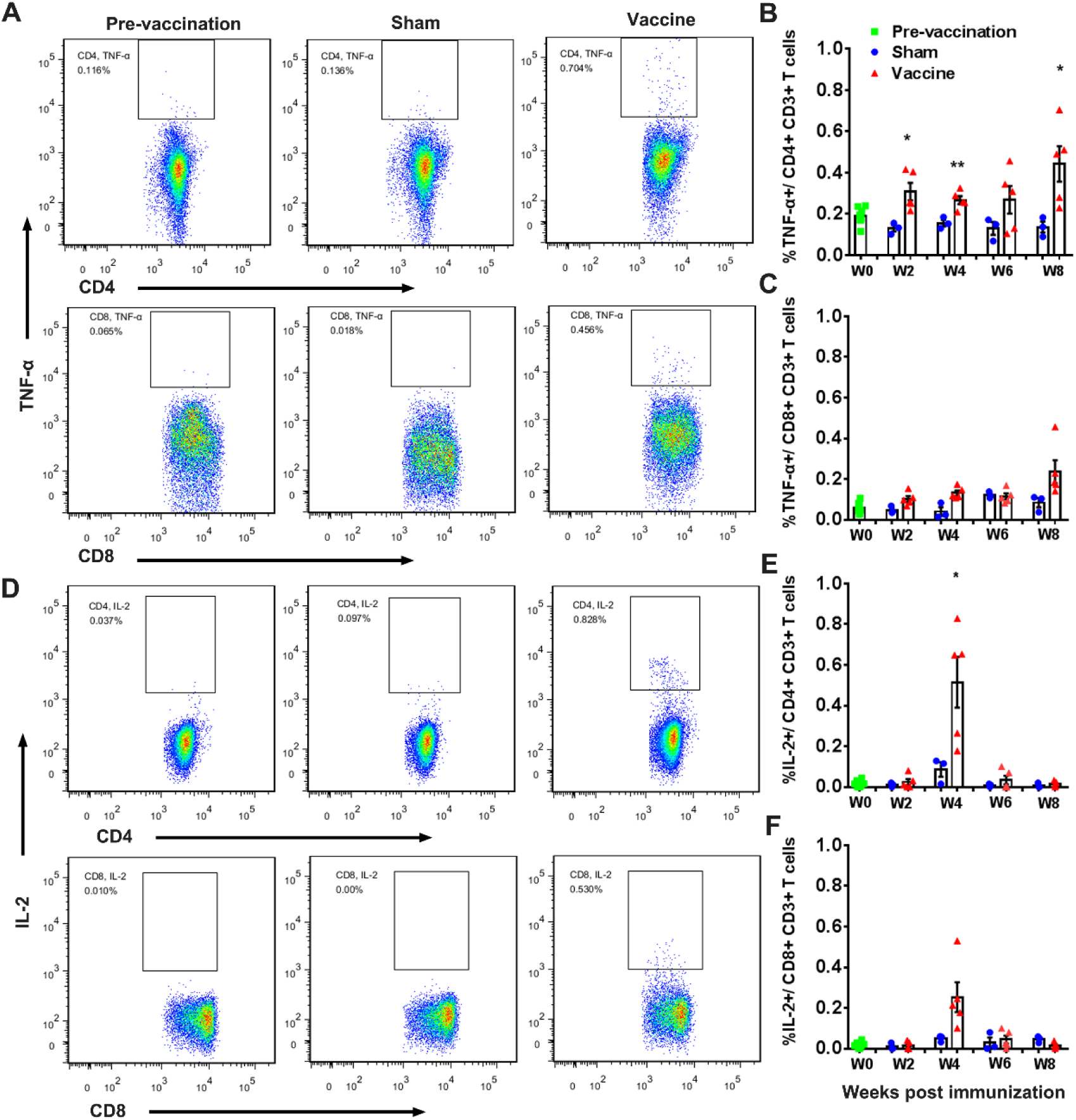
Frequency of intracellular TNFα and IL-2 expressing T cell response in PBMCs from rhesus macaques immunized with Sad23L-nCoV-S and Ad49L-nCoV-S vaccines or sham controls. (A-C) Frequency of TNF-α^+^ CD4^+^/CD8^+^ T cell response to S peptides. (D-F) Frequency of TNF-α^+^ CD4^+^/CD8^+^ T cell response to S peptides. Data were shown as means ± SEM (standard errors of means). *P* values were analyzed by Student’s *t* test. Statically significant differences were showed with asterisks (*, *P*<0.05; **, *P*< 0.01 and ***, *P*< 0.001). ns, *P*>0.05 and no significant difference.

**Fig. S8.**
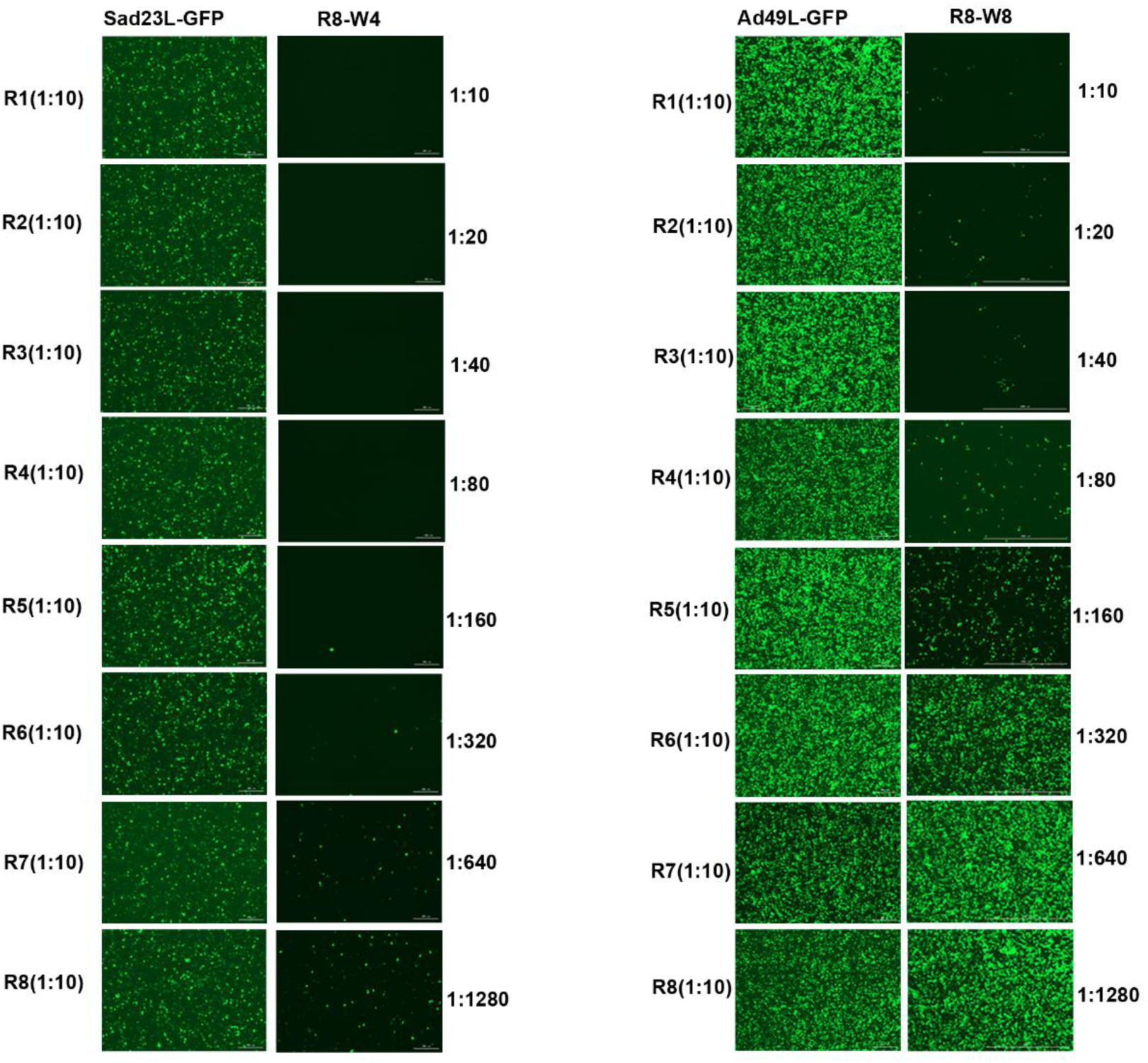
Measurement of serum NAb titers to Ad49L and Sad23L vectors in vaccinated macaques. Two representative plasma samples of vaccinated macaque (R8 at weeks 4 and 8) were tested on HEK-293A cells for neutralizing Sad23L-GFP and Ad49L-GFP vectorial viruses by green fluorescent activity assay. Neutralization titer was defined as the maximum serum dilution that neutralized 50% of Green fluorescent activity.

**Fig. S9.**
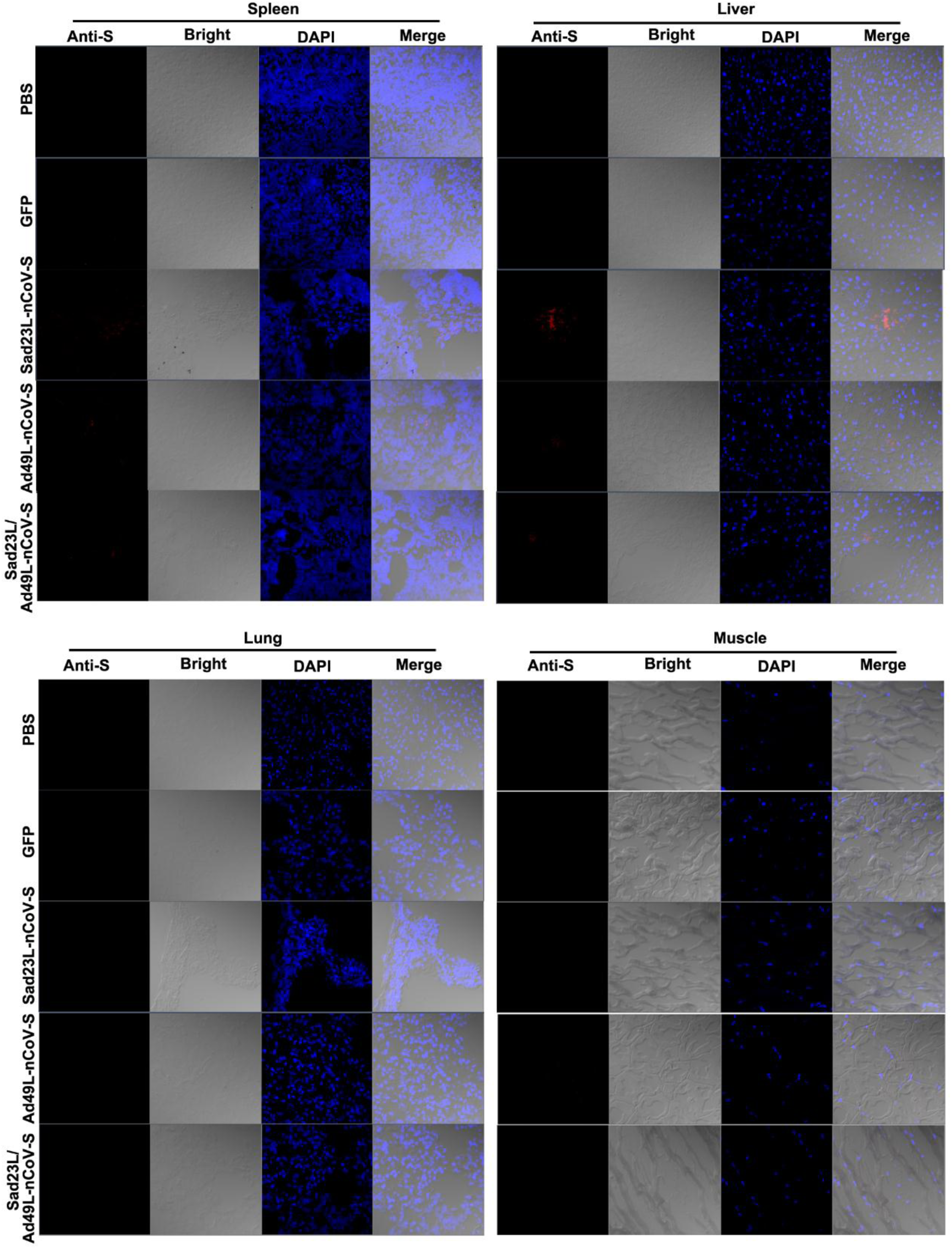
Examination of SARS-CoV-2 S protein in the tissues of Sad23L-nCoV-S and Ad49L-nCoV-S immunized mice. Spleen, liver, Lung and muscle tissues of immunized C57BL/6 mice were examined by immunofluorescence staining with a human monoclonal antibody to SARS-CoV-2 S and DAPI.

**Table S1.** Basic information of rhesus macaques pre-vaccination

| Group | ID | Macaques | Age | Weight | NAb titer of pre-exposure to |  |  |
| --- | --- | --- | --- | --- | --- | --- | --- |
|  |  | (gender) | (year) | (kg) | Ad5 | Sad23L | Ad49L |
| Sham | 00159 | R3(M) | 13 | 11.79 | <1:10 | <1:10 | <1:10 |
|  | 06120011 | R4(M) | 13 | 6.76 | <1:10 | <1:10 | <1:10 |
|  | 08050591 | R6(M) | 12 | 7.99 | <1:10 | <1:10 | <1:10 |
| Vaccination | 01191 | R1 (M) | 14 | 11.56 | <1:10 | <1:10 | <1:10 |
|  | 01399 | R2(M) | 13 | 7.51 | <1:10 | <1:10 | <1:10 |
|  | 00943 | R5(M) | 13 | 9.32 | <1:10 | <1:10 | <1:10 |
|  | 00807 | R7(M) | 12 | 10.81 | <1:10 | <1:10 | <1:10 |
|  | 08040661 | R8 (M) | 11 | 8.72 | <1:10 | <1:10 | <1:10 |

**Table S2.** Measuring of hematological and biochemistry indexes of rhesus macaques in the course of pre- and post-vaccination with Sad23L-nCoV-S and Ad49L-nCoV-S vaccines

| Animal<br>Days |  | R1 | R2 | R3 | R4 | R5 | R6 | R7 | R8 |
| --- | --- | --- | --- | --- | --- | --- | --- | --- | --- |
| TP | 0 | 71.12 | 69.8 | 72.78 | 74.1 | 71.96 | 72.16 | 68.78 | 76.89 |
|  | 7 | 64.37 | 67.35 | 73.1 | 69.97 | 70.21 | 69.94 | 66.56 | 71.74 |
|  | 21 | 72.1 | 73.03 | 76.25 | 72.76 | 76.67 | 73.23 | 72.38 | 75.28 |
|  | 42 | 71.32 | 76.57 | 75.93 | 72.75 | 79.37 | 77.27 | 73.05 | 76.59 |
|  | 56 | 75.88 | 70.74 | 74.55 | 73.72 | 78.55 | 73.88 | 72.9 | 74.13 |
| Alb | 0 | 43.74 | 45.41 | 42.93 | 44.01 | 40.7 | 44.72 | 44.71 | 45.38 |
|  | 7 | 39.68 | 41.73 | 43.36 | 43.43 | 40.82 | 41.59 | 42.44 | 41.89 |
|  | 21 | 44.74 | 46.31 | 47.16 | 44.55 | 46.3 | 46.27 | 46.18 | 44.97 |
|  | 42 | 43.41 | 47.02 | 45.47 | 42.23 | 44.72 | 45.54 | 45.25 | 44.64 |
|  | 56 | 46.77 | 44.79 | 46.59 | 43.68 | 46.81 | 46.19 | 47.98 | 43.87 |
| Glb | 0 | 27.38 | 24.39 | 29.85 | 30.09 | 31.26 | 27.44 | 24.07 | 31.51 |
|  | 7 | 24.69 | 25.62 | 29.74 | 26.54 | 29.39 | 28.35 | 24.12 | 29.85 |
|  | 21 | 27.36 | 26.72 | 29.09 | 28.21 | 30.37 | 26.96 | 26.2 | 30.31 |
|  | 42 | 27.91 | 29.55 | 30.46 | 30.52 | 34.65 | 31.73 | 27.8 | 31.95 |
|  | 56 | 29.11 | 25.95 | 27.96 | 30.04 | 31.74 | 27.69 | 24.92 | 30.26 |
| A/G | 0 | 1.6 | 1.86 | 1.44 | 1.46 | 1.3 | 1.63 | 1.86 | 1.44 |
|  | 7 | 1.61 | 1.63 | 1.46 | 1.64 | 1.39 | 1.47 | 1.76 | 1.4 |
|  | 21 | 1.64 | 1.73 | 1.62 | 1.58 | 1.52 | 1.72 | 1.76 | 1.48 |
|  | 42 | 1.56 | 1.59 | 1.49 | 1.38 | 1.29 | 1.44 | 1.63 | 1.4 |
|  | 56 | 1.61 | 1.73 | 1.67 | 1.45 | 1.47 | 1.67 | 1.93 | 1.45 |
| TBil | 0 | 5.85 | 3.65 | 2.97 | 3.1 | 2.93 | 3.56 | 4.76 | 3.59 |
|  | 7 | 3.32 | 1.81 | 2.39 | 3.17 | 1.8 | 5.88 | 3.41 | 3.04 |
|  | 21 | 1.95 | 2.42 | 2.93 | 2.7 | 2.46 | 3.3 | 1.58 | 3.19 |
|  | 42 | 1.91 | 1.31 | 1.8 | 1.33 | 1.67 | 1.66 | 1.78 | 1.2 |
|  | 56 | 2.21 | 1.22 | 1.8 | 1.6 | 2.28 | 1.55 | 1.38 | 2.38 |
| ALT | 0 | 100 | 57.7 | 61.3 | 91.8 | 47.7 | 69.7 | 138.6 | 83.8 |
|  | 7 | 96.8 | 65.5 | 44.5 | 61.1 | 55.4 | 78 | 103.3 | 121 |
|  | 21 | 65.9 | 53.9 | 56.4 | 32.6 | 62.2 | 36.9 | 76.3 | 96.8 |
|  | 42 | 63.4 | 46.4 | 70.9 | 21.8 | 63.7 | 36.4 | 56.7 | 49.3 |
|  | 56 | 39.9 | 33.3 | 49 | 19.4 | 48.1 | 24.5 | 52.2 | 27.7 |
| AST | 0 | 49.7 | 36.7 | 28.8 | 60.2 | 49.7 | 26.2 | 77.3 | 39.1 |
|  | 7 | 47.5 | 33.3 | 32.5 | 51 | 37.9 | 28.9 | 56.3 | 42.4 |
|  | 21 | 31.7 | 29.8 | 25 | 32.7 | 34.8 | 21.2 | 42.3 | 50.1 |
|  | 42 | 27.4 | 31.6 | 31.6 | 29.8 | 33.1 | 21.3 | 31.9 | 28.4 |
|  | 56 | 28.5 | 26.2 | 29 | 33.5 | 31.7 | 20.5 | 36.9 | 26.5 |
| **GGT**<br>**U/L** | 0 | 97.9 | 42.7 | 62.7 | 57.6 | 89.2 | 140.5 | 54.5 | 68.3 |
|  | 7 | 88 | 50.1 | 64.3 | 56.3 | 74.5 | 128.8 | 75.9 | 76 |
|  | 21 | 110.9 | 71.8 | 83.2 | 55 | 77.7 | 164 | 64.9 | 83.5 |
|  | 42 | 84.6 | 74.2 | 77.6 | 52.2 | 70.8 | 169.4 | 56.6 | 77.8 |
|  | 56 | 78.2 | 60.6 | 63.3 | 48.3 | 64.7 | 125 | 61.8 | 70.1 |
| **ALP**<br>**U/L** | 0 | 95.85 | 141.27 | 105.9 | 201.57 | 85.65 | 123.09 | 106.01 | 184.39 |
|  | 7 | 110.17 | 265.49 | 118.37 | 229.04 | 117.7 | 179.86 | 142.02 | 193.16 |
|  | 21 | 113.51 | 220.7 | 119.59 | 183.41 | 83.99 | 149.54 | 116.65 | 161.14 |
|  | 42 | 112.95 | 217.72 | 91.87 | 151.17 | 63.11 | 106.35 | 83.32 | 142.87 |
|  | 56 | 90.59 | 168.74 | 88.32 | 122.93 | 50.26 | 77.8 | 86.85 | 142.28 |
| **CK**<br>**U/L** | 0 | 153.4 | 122.7 | 138.3 | 166.7 | 168.6 | 118.5 | 245.5 | 81.4 |
|  | 7 | 97.8 | 156.8 | 87 | 165.1 | 149.2 | 129.6 | 233.5 | 70.1 |
|  | 21 | 84.1 | 139 | 103.1 | 118.9 | 145.3 | 140.5 | 111.7 | 76.7 |
|  | 42 | 75.6 | 136.6 | 86 | 93 | 163.9 | 81.3 | 128 | 64.9 |
|  | 56 | 83.6 | 101.9 | 101.1 | 90.9 | 107.1 | 71.5 | 108.2 | 53 |
| **LDH**<br>**U/L** | 0 | 344.2 | 302.9 | 215.7 | 336.3 | 275.3 | 224.2 | 430.9 | 198.9 |
|  | 7 | 368.7 | 265.6 | 237.2 | 378.8 | 233.4 | 331.6 | 467.2 | 205.6 |
|  | 21 | 220.9 | 219.2 | 170.1 | 187.3 | 209.2 | 180.1 | 316.6 | 209.6 |
|  | 42 | 199.4 | 253.9 | 155 | 188.5 | 231.2 | 217.7 | 266 | 189.9 |
|  | 56 | 210.4 | 241.1 | 202.5 | 185.8 | 235 | 216.5 | 365.4 | 164.7 |
| **GLU**<br>**mmol/L** | 0 | 5.63 | 4.79 | 5.88 | 6.2 | 4.94 | 6.66 | 5.11 | 4.59 |
|  | 7 | 3.9 | 3.81 | 4.75 | 4.87 | 4.34 | 4.59 | 3.58 | 4.6 |
|  | 21 | 4.67 | 4.66 | 5.25 | 5.75 | 5.24 | 4.58 | 5.57 | 5 |
|  | 42 | 4.17 | 4.52 | 4.79 | 5.46 | 6.19 | 6.22 | 5.31 | 5.42 |
|  | 56 | 4.14 | 4 | 4.83 | 5.56 | 4.2 | 6.23 | 4.85 | 4.87 |
| **BUN**<br>**mmol/L** | 0 | 7.5 | 7.3 | 3.9 | 7.5 | 7.1 | 8.6 | 7.8 | 6.7 |
|  | 7 | 8.3 | 5.3 | 5.5 | 10.3 | 5.8 | 5.1 | 7.1 | 6.1 |
|  | 21 | 9.1 | 7.1 | 5.3 | 5.2 | 7.1 | 7.3 | 7.8 | 6.4 |
|  | 42 | 6.4 | 6.5 | 6.7 | 7.1 | 5.8 | 6.2 | 7.9 | 6.6 |
|  | 56 | 7.1 | 6.1 | 5.2 | 4.6 | 5.8 | 6.6 | 7.6 | 6.1 |
| **CREA**<br>**μmol/L** | 0 | 114.3 | 74.9 | 76.6 | 84.7 | 74.2 | 85.4 | 82.9 | 71.4 |
|  | 7 | 95.9 | 65.9 | 77.8 | 90.1 | 82.1 | 77 | 83.6 | 66.8 |
|  | 21 | 103.3 | 66.5 | 83.7 | 75.2 | 83.6 | 81.1 | 84 | 66.5 |
|  | 42 | 97.3 | 62.6 | 80.5 | 75.2 | 76.6 | 72.1 | 71.4 | 59.9 |
|  | 56 | 107.5 | 53 | 78 | 68.9 | 78.9 | 71.6 | 80.8 | 57.7 |
| **CHOL**<br>**mmol/L** | 0 | 2.82 | 2.49 | 2.34 | 3.16 | 2.81 | 2.87 | 2.33 | 3.05 |
|  | 7 | 2.7 | 3.11 | 2.7 | 3.47 | 2.74 | 3.2 | 2.51 | 3.16 |
|  | 21 | 2.47 | 2.9 | 3.05 | 3.53 | 2.78 | 3.57 | 2.37 | 3.17 |
|  | 42 | 2.6 | 3.35 | 3.01 | 3.46 | 3.12 | 3.77 | 2.65 | 3.37 |
|  | 56 | 3.22 | 3.05 | 3.26 | 3.73 | 3.02 | 3.91 | 2.97 | 3.71 |
| **TG** | 0 | 0.17 | 0.34 | 0.43 | 0.5 | 0.86 | 0.52 | 0.45 | 0.35 |
|  | 7 | 0.18 | 0.4 | 0.53 | 0.36 | 0.73 | 0.71 | 0.35 | 0.32 |
| **mmol/L** | 21 | 0.27 | 0.7 | 0.69 | 0.66 | 0.65 | 0.61 | 0.52 | 0.38 |
|  | 42 | 0.24 | 0.54 | 0.65 | 0.66 | 0.65 | 0.55 | 0.62 | 0.56 |
|  | 56 | 0.33 | 0.54 | 0.55 | 0.84 | 0.53 | 1.08 | 0.54 | 0.85 |

**Table S3.**
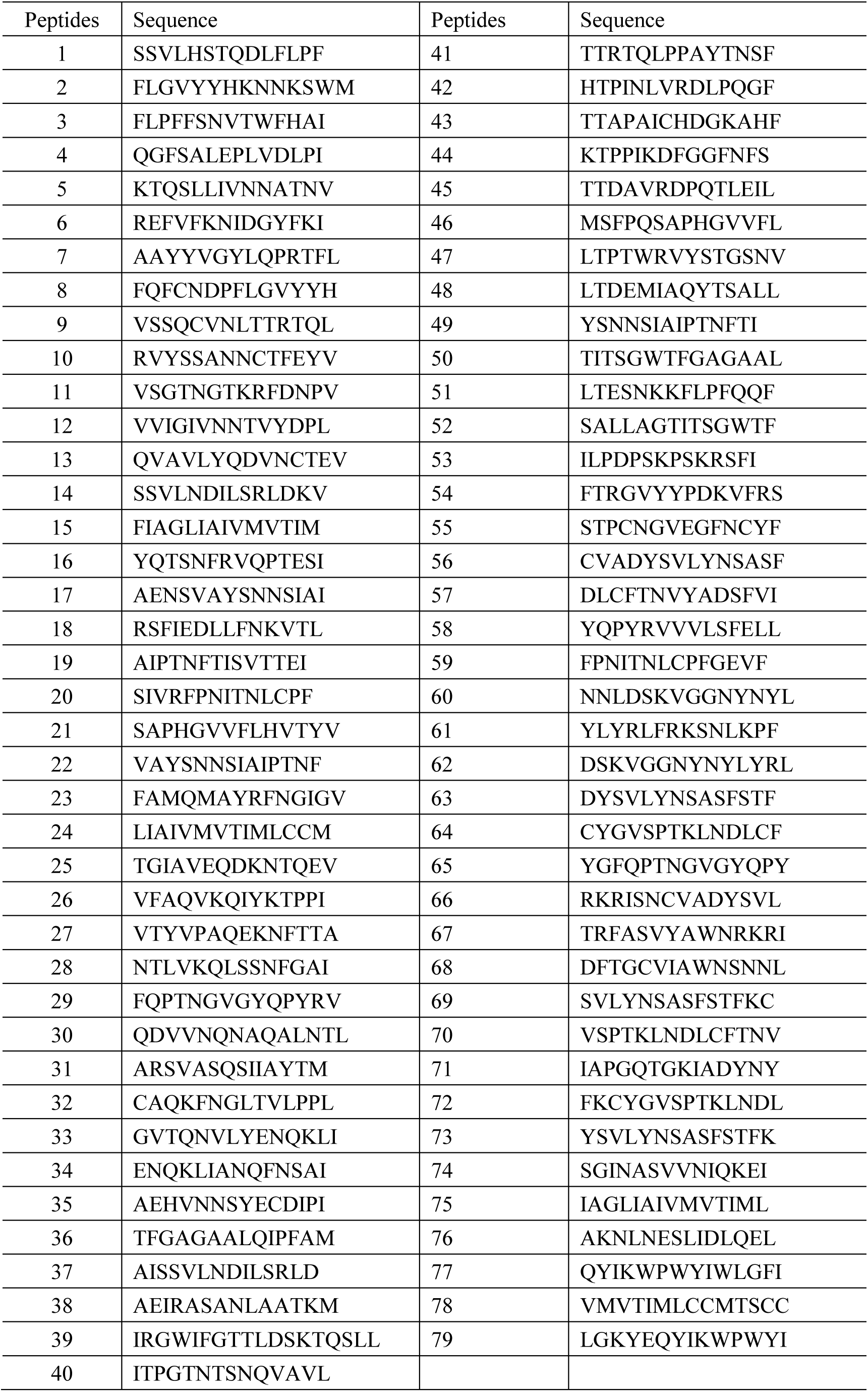
Peptides derived from amino acid sequences of SARS-CoV-2 S protein used in ELISpot and ICS

**Table S4.**
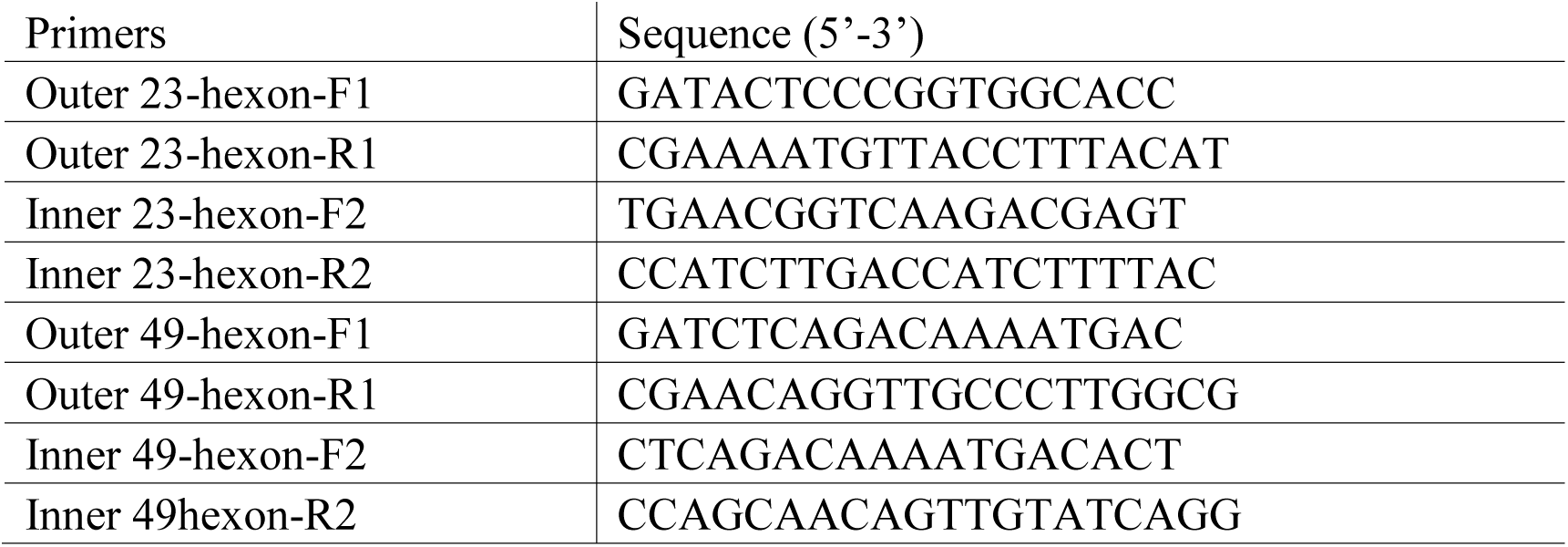
Nested-PCR primers specific for hexon of Sad23L or Ad49L vector

